# Purifying selection on noncoding deletions of human regulatory elements detected using their cellular pleiotropy

**DOI:** 10.1101/2020.05.19.105205

**Authors:** David W. Radke, Jae Hoon Sul, Daniel J. Balick, Sebastian Akle, Alzheimer’s Disease Neuroimaging Initiative, Robert C. Green, Shamil R. Sunyaev

## Abstract

Genomic deletions provide a powerful loss-of-function model in non-coding regions to assess the role of purifying selection on human noncoding genetic variation. Regulatory element function is characterized by non-uniform tissue/cell-type activity, necessarily linking the study of fitness consequences from regulatory variants to their corresponding cellular activity. We used deletions from the 1000 Genomes Project (1000GP) and a callset we generated from genomes of participants in the Alzheimer’s Disease Neuroimaging Initiative (ADNI) in order to examine whether purifying selection preserves noncoding sites of chromatin accessibility (DHS), histone modification (enhancer, transcribed, polycomb-repressed, heterochromatin), and topologically associated domain loops (TAD-loops). To examine this in a cellular activity-aware manner, we developed a statistical method, Pleiotropy Ratio Score (PlyRS), which calculates a correlation-adjusted count of “cellular pleiotropy” for each noncoding base-pair by analyzing shared regulatory annotations across tissues/cell-types. Comparing real deletion PlyRS values to simulations in a length-matched framework and using genomic covariates in analyses, we found that purifying selection acts to preserve both DHS and enhancer sites, as evident by both depletion of deletions overlapping these annotations and a shift in the allele frequency spectrum of overlapping deletions towards rare alleles. However, we did not find evidence of purifying selection for transcribed, polycomb-repressed, or heterochromatin sites. Additionally, we found evidence that purifying selection is acting on TAD-loop boundary integrity by preserving co-localized CTCF binding sites. Notably, at regions of DHS, enhancer, and CTCF within TAD-loop boundaries we found evidence that both sites of tissue/cell-type-specific activity and sites of cellularly pleiotropic activity are preserved by selection.

**Significance Statement:** We used natural genomic deletions as a loss-of-function model to assess the role of purifying selection in preserving human noncoding regulatory sites. We examined this in a cellular activity-aware manner through development of a statistical method, Pleiotropy Ratio Score (PlyRS), which calculates an adjusted count of “cellular pleiotropy” for each noncoding basepair by analyzing correlations from shared regulatory annotations across tissues/cell-types. By comparing real deletion PlyRS values to simulations, we found that purifying selection acts to preserve both DHS and enhancer sites and TAD-loop boundary integrity by preserving co-localized CTCF binding sites. Notably, we found evidence at these regulatory regions that both sites of tissue/cell-type-specific activity and sites of cellularly pleiotropic activity are preserved by selection.

---

**L**arge-scale sequencing studies have provided tremendous insight into biological function and human disease, with statistical signatures of natural selection serving as a primary identifying feature. The classic example is the analysis of selective constraints on protein coding genes evident from the depletion of missense or nonsense genetic variants. These advances, however, are not directly translatable to the analysis of noncoding DNA, which has increasingly become a focus of human genetics research. Functional genomic studies have revealed numerous regions of regulatory activity marked by chromatin accessibility and histone modification (1, 2). Association signals for common human phenotypes are dramatically enriched in these regulatory regions of the genome (3), showcasing the importance of specialized cellular function. In contrast to protein-coding sequences, the function of regulatory sequences is not determined by triplet codon structure thereby providing no obvious analog to protein-truncating single nucleotide variants (SNVs) to identify loss of function. This ambiguity of the mutational consequences of individual nucleotides within regulatory sequences complicates the ability to study their function through the lens of purifying natural selection. Previous work focusing on SNVs within noncoding regions developed sophisticated genetic models that relied on functional proxies such as transcription factor binding sites, nucleotide conservation across species, or machine learning (4–11). However, it is difficult to clearly interpret these findings in terms of selection against the loss of regulation. In contrast to SNVs, deletions are a class of variation that provide a direct loss of normal regulatory function at a locus by physically removing the sequence of a regulatory element in at least a heterozygous manner. This logic underlies experimental studies of regulatory function using CRISPR/Cas9 systems (12, 13). Yet, natural population genetic variation provides a more systematic and genome-wide view of the action of selection on deletions. Work done by sequencing consortia has demonstrated reduction of deletion variation in various categories of regulatory sequences (14–16).

The hallmark feature of human regulatory elements is their non-uniform activity across tissues and cell types. Here, we offer a population genetic analysis of natural deletions in light of variable regulatory activity across tissues. Deletions that remove sites of genomic regulation with pleiotropic cellular effects (what we term “cellular pleiotropy”, i.e. the same regulatory element locus is active in more than one tissue or cell-type) might be expected to be, on average, more deleterious (i.e. fitness-reducing) than deletions that remove cell-type-specific sites, since any changes at the DNA level to these regulators potentially affects multiple tissues/cell-types simultaneously. Another possibility is that since tissue/cell-type-specific regulation is what enables widespread cellular diversity, these regulatory elements must be under strong selective constraint to preserve their specialized biological function. These two potential modes of selection preserving regulation of cellular activity are not mutually exclusive, as selection may be operating to remove overlapping deletions to preserve the utility of both types of regulators. Prior work has provided suggestive evidence that tissue activity count is a contributor to selective constraint in regulatory sequences (10, 16, 17). Studying purifying selection on noncoding deletions is thus inherently tied to the cellular activity of corresponding deleted regulatory sequences. To address this, we have developed a statistical method, Pleiotropy Ratio Score (PlyRS), to quantify the amount of tissue/cell-type activity (i.e. cellular pleiotropy) for individual nucleotides in light of the hierarchical developmental structure of human tissues and cell types, while controlling for their correlation rather than using a simple tissue/cell-type count. We then analyzed separately several diverse epigenomic features (open chromatin, histone modifications, and topologically associated domain loops) taking into account non-independence of these individual annotations across tissues and cell types using our PlyRS values. In this way, we assessed the effect of purifying selection on millions of nucleotide positions in the human genome by examining patterns of PlyRS values within naturally occurring deletion sequences.

Reduction of genetic variation and a shift in the allele frequency spectrum (AFS) towards rare variants are two key signatures of purifying selection. If selection is operating on the removal of deleterious deletions overlapping regulatory regions, we would expect to see both a reduction in deletion variation overlapping the important regulatory features and a shift in the AFS of remaining overlapping deletions towards rarer frequencies, relative to neutral expectations. These conditions on segregating deletions should be simultaneously present to conclude that purifying natural selection is acting to preserve a particular regulatory epigenomic feature(s), as either reduced deletion counts or a shift in the deletion AFS alone may indicate deletion calling artifacts or confounding genomic covariates. Both of these signatures are prone to various biological and technical confounders, particularly for structural variation. For example, the accuracy of deletion calls is influenced by their length and allele frequency (AF) (18). Longer deletions have more prevalent missing coverage and common deletions are observed more often in the population, so these types of deletions are more likely to be correctly identified using current methods based on analyzing short-read sequencing data. Variant calling accuracy also depends on the mappability of the sequence (19). Another known issue is the observed negative correlation of deletion length and AF (14, 20). This could be due to underlying biology, deletion caller algorithm biases, or both. In addition to technical confounders, biological factors unrelated to the direct pressure of selection may affect the degree of variation (i.e. the number of segregating mutations) and the AFS. For example, the degree of variation is linearly proportional to mutation rate; however, the deletion mutation rate at fine-scale is still unknown and could be influenced by sequence GC content and other local genomic properties. In contrast to the overall variation, the normalized AFS is not affected by mutation rate, at least for relatively small sample sizes, but together with degree of variation could be influenced by complex mechanisms like background selection. To address these complications, we simulated length-matched positions of each real deletion while keeping the original AF label, and took into account relevant genomic confounding variables co-occurring with the same deletion. Using this framework, we compared the observed diversity and AFS of real deletions to the expectations based on computer simulations using analyses of PlyRS values across their coordinates.

## Results

### Pleiotropy Ratio Score (PlyRS)

To score deletions with respect to their effect on regulatory function, we considered both the number of removed elements and the activity of each element across cell and tissue types. In contrast to SNVs, a noncoding deletion can potentially remove regulatory function at a genomic locus along two distinct “axes” (see SI Appendix, Fig. S1 for a cartoon). One axis (“horizontal”) corresponds to the amount of regulatory space removed by the deletion irrespective of its tissue/cell-type activity. The other axis (“vertical”) corresponds to the combined amount of regulatory activity across tissues and cell types of each base-pair (i.e. the cellular pleiotropy of a regulatory coordinate). Thus, for any deletion overlapping regulatory sequences, there will be a simultaneous removal at that locus along both axes, which we quantify by a counting score for each axis.

For the horizontal axis we count deleted base-pairs with a regulatory annotation from any tissue/cell-type (*SI Appendix, Note S1a*). We do not require removal of an entire regulatory element for this horizontal count, since deletion of even a partial regulatory element sequence can render it inoperable (21). Additionally, since regulatory element boundaries are not perfectly aligned between tissues, it could be the case that a partial deletion of an element observed in one tissue may correspond to a complete deletion of the element observed in another tissue. Consequently, for a deletion overlapping a regulatory element(s), the horizontal axis count score can range from as low as 1 (only a single regulatory base-pair deleted) to as high as the length of the deletion (all base-pairs along the deletion length overlap a regulatory element[s]).

A simple numerical count of the number of tissues/cell-types where a regulatory element locus has activity is not sufficient for properly specifying cellular pleiotropy, because this count can be heavily influenced by the cellular diversity of the particular tissues/cell-types included in the analysis. For example, a count of 3 in an analysis performed with heart tissue, lung tissue, and ten blood cell-types would not have the same interpretation as a count of 3 in an analysis performed with heart tissue, lung tissue, and only one blood cell-type. In the former, it could be that the count of 3 comes from three highly-correlated blood cell-types, but in the latter, the count of 3 would have to come from the more developmentally diverse set of all three tissues/cell-types. Therefore, to enable proper “counting” of cellular pleiotropy, we developed a statistical method, called Pleiotropy Ratio Score (PlyRS), which calculates a correlation-adjusted count of cellular pleiotropy for each base-pair in the noncoding genome (*SI Appendix, Note S1b*).

We use the PlyRS value at any given base-pair along the length of a deletion to provide the counting score along the vertical axis. At any base-pair coordinate within a deletion, the PlyRS value can range from 0-indicating no tissue/cell-type included in the analysis has annotated activity-to a maximum of 1-indicating that all tissues/cell-types analyzed have annotated activity. Between these extreme bounds, PlyRS does not simply calculate the fraction of tissues/cell-types where the base-pair exhibits regulatory activity, rather it weights this proportion relative to the overall correlation of regulatory activity across these tissues/cell-types along the genome. For example, a base-pair active in three highly related cell types would be assigned a lower PlyRS value than a base-pair active in three (or potentially less) unrelated cell types. Thus, as a consequence of the PlyRS method calculation, counts of elements active in tissues/cell-types with common activity will be down-weighted while counts of elements active in tissues/cell-types with rare activity will be up-weighted. Similarly, for each base-pair that has only tissue/cell-type-specific activity, the PlyRS value will be different for that particular tissue/cell-type depending on how its activity covaries across the genome with the other regulatory tissues/cell-types being analyzed. *SI Appendix* Fig. S2 and Fig. S3, respectively, illustrate how PlyRS corresponds to the raw tissue/cell-type count and how PlyRS compares to tissue/cell-type-specific counts.

### Construction of Deletion and Regulatory Datasets

To examine potential selective constraints on deletions within regulatory regions, we needed fine-resolution of genomic coordinates for both deletions and regulatory regions as well as high-confidence deletion allele frequencies from population data. For this, we compiled deletion data from two callsets and regulatory data from seven callsets, and applied additional filters relevant to our analysis. See Materials and Methods for additional criteria used to ensure high-quality datasets.

We used deletion data from the 1000 Genomes Project Consortium Phase 3 callset (1000GP) of breakpoint-resolved deletions for which deletions were genotyped in 2,504 individuals from 26 modern human populations (14) (*SI Appendix, Note S2a*). We additionally used deletions that we called and genotyped across 752 individuals sequenced as part of the Alzheimer’s Disease Neuroimaging Initiative (ADNI) (22) (*SI Appendix, Note S2b* and *Note S3*), using the CNV algorithm GenomeSTRiP (23). We restricted our analysis to noncoding deletions. As expected, the bulk (>80%) of deletions in our datasets remaining after filtering were rare (below 1% AF).

To analyze genomic deletions within regulatory regions, we used regulatory data from the NIH Roadmap Epigenomics Consortium (REC) (2). In particular, we used two callsets of chromatin accessibility data (DNase I hypersensitivity “DHS”) and four callsets of histone modification data (H3K4me1 “enhancer”, H3K36me3 “transcribed”, H3K27me3 “polycomb-repressed”, and H3K9me3 “heterochromatin”). Two sets of DHS annotation (hotspot and MACS) were used to check for consistency in the analyses. DHS annotations are typically associated with sites of open chromatin allowing accessibility for regulator binding and histone annotations are typically associated with sites of specific regulatory activity, as noted. We additionally used regulatory data that demarcate topologically associated domain loops (TAD-loops) (24), which are associated with local genomic regions of physically interacting regulatory activity.

### Depletion of Variation at DHS or Enhancer Sites

We first tested whether there was evidence of depletion of noncoding deletion variation overlapping chromatin accessibility and histone modifications (*SI Appendix, Note S4a*). We corrected for the confounding effects of mappability, deletion length, and allele frequency using simulations. For each real deletion in both the 1000GP and ADNI datasets, we randomly simulated 1,000 deletions of the same length to occur on the same chromosome and same noncoding genomic compartment space (intronic or intergenic) using only uniquely mappable sequence coordinates (*SI Appendix, Note S2d*) for both real deletions and simulated deletions. For more detail on our length-matched simulations, see *SI Appendix, Notes S5a-S5c*. We summed the PlyRS values calculated per base-pair along the length of every deletion. This sum, denoted PlyRS_sum_ (*SI Appendix, Note S1c*), corresponds to the total cellular pleiotropy (for a specific regulatory feature) of the deletion, encompassing both the horizontal and vertical “axes” along which purifying selection may be operating on the deletion (*SI Appendix, Note S1*). We compared PlyRS_sum_ values for both real and simulated deletions and quantified depletion across regulatory features using Cohen’s D statistic (*SI Appendix, Note S5b*).

Panel A of Fig. 1 shows PlyRS_sum_ effect sizes from comparing real data to simulations and indicates significant depletion of deletions (Cohen’s D>2, corresponding to 2 std. dev.) overlapping DHS or enhancer regions. We did not detect a significant depletion for deletions overlapping transcribed, polycomb-repressed, or heterochromatin epigenomic features. The depletion of deletions overlapping DHS or enhancer sites was significant not only in the full deletion sets, but also in both the intronic and intergenic genomic compartments. Additionally, we found concordance between effect sizes in 1000GP and ADNI datasets for DHS or enhancer deletion depletions, suggesting reliable capture of biological information from deletion callsets with differing characteristics (*SI Appendix*, Tables S1-S2). These results suggest that purifying selection may be operating broadly on deletions to preserve DHS and enhancer epigenomic features. *SI Appendix*, Table S5d1 (*Note S5d*) lists the effect sizes found in the depletion simulations.

**Fig. 1.**
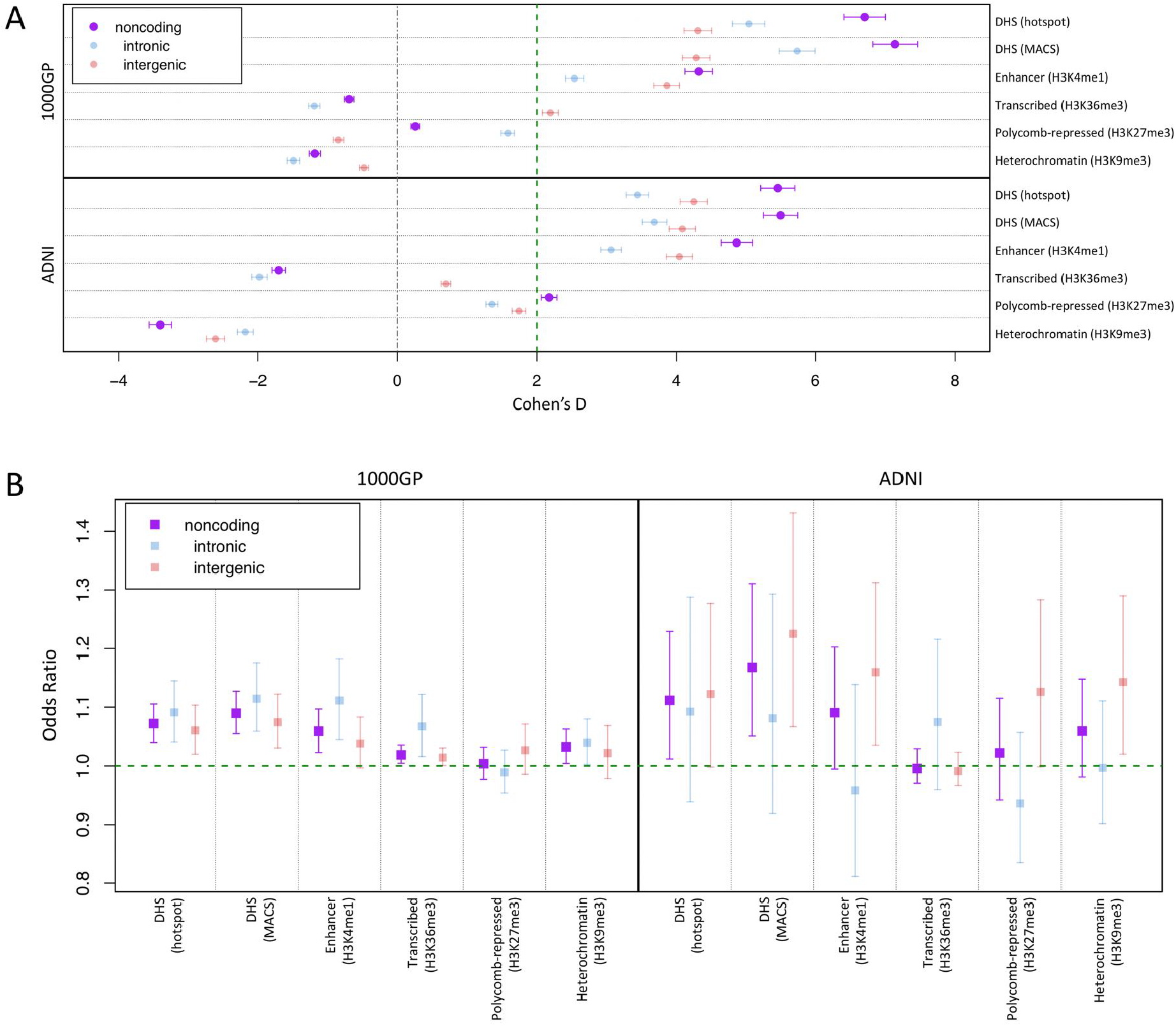
Depletion of deletions and shift of deletion allele frequency spectrum overlapping regulatory sites. (A) We calculated PlyRS_sum_ (*SI Appendix, Note S1c*) for every deletion to quantify overlap with sites of chromatin accessibility or histone modification. We plot the degree of reduction in the PlyRS_sum_ for real deletions relative to simulation. This reduction is measured using Cohen’s D, which is the effect size of a t-test on PlyRS_sum_ values (*SI Appendix, Notes S5a-S5b*) in units of standard deviation (plotted with 95% confidence intervals on the mean reduction). Two units of effect size (Cohen’s = 2) approximately corresponds to the 95% confidence interval of significance in depletion. Higher values of Cohen’s D indicate larger depletion within those sets compared to simulation. In presence of the true effect, there is sample size dependence on the underlying t-test, and the expected value of Cohen’s D would be higher for larger datasets. (B) For each deletion we determined the magnitude of PlyRS_sum_ depletion, calculated as a ratio between its PlyRS_sum_ and the average PlyRS_sum_ of its length-matched simulated deletions (*SI Appendix, Note S6b*), for sites of chromatin accessibility or histone modification. We tested whether PlyRS_sum_ depletion magnitude depends on allele frequency (deletions categorized as rare [AF<=1%] or common), using multivariate logistic regression in the presence of genomic covariates (*SI Appendix, Note S6a*). We plot the regression odds ratio (OR) with 95% profile likelihood-based confidence intervals. Results above 1 indicate positive correlation of the magnitude of PlyRS_sum_ depletion with allele frequency. This corresponds to an excess of rare alleles overlapping the regulatory feature in the real dataset compared to simulation, which is the expected result for features being preserved by the action of purifying selection against overlapping deletions.

### Shift in Allele Frequency Spectrum at DHS or Enhancer Sites

We next tested whether there was a shift in the allele frequency spectrum of noncoding deletions overlapping the chromatin accessibility and histone modification epigenomic features. The analysis of allele frequency distribution is important because the total degree of variation can be confounded by mutation rate (unlike SNVs, we do not have good models for mutation rate along the genome for deletions [(25)]). The allele frequency distribution, when normalized, does not depend on mutation rate for relatively small populations (within the limits of the infinite sites approximation), but due to the recent explosive growth of the human population, this assumption may break down for extremely large sample sizes at which point recent recurrent mutations become relevant. However, for the sample sizes analyzed here, the allele frequency distribution can be assumed to be independent of mutation rate, with the chance of recurrent mutations being small. This is especially true for deletions which would require recurrent mutations to occur at the same breakpoints (start and end coordinates being identical). Therefore, a shift in the allele frequency spectrum of real deletions in our datasets compared to simulated deletions would likely reflect the action of purifying selection. Still, the allele frequency distribution can be affected by a number of variables unrelated to selective pressure. To take into account the potential effect of background selection, we controlled for regional (50kb +/- deletion coordinates) SNV nucleotide diversity and recombination rate, as well as distance to the nearest transcription start site. We additionally controlled for regional GC content. Due to technical confounders, allele frequency is expected to be influenced by deletion length so we also controlled for length explicitly. We accounted for these genomic covariates using multivariate logistic regression, testing whether PlyRS_sum_ depletion magnitude depended on allele frequency (deletions categorized as rare [AF<=1%] or common; *SI Appendix, Note S6a*). To measure the magnitude of potential PlyRS_sum_ depletion for each deletion, we calculated a ratio between its PlyRS_sum_ and the average PlyRS_sum_ of its length-matched simulated deletions (*SI Appendix, Note S6b*). If purifying selection is, in fact, acting against deletions overlapping regulatory features, we would expect the largest PlyRS_sum_ depletions to be found in common deletions (in our test, an odds ratio [OR] above 1 which shows positive correlation with allele frequency).

Panel B of Fig. 1 shows that for deletions overlapping DHS sites the OR significantly (confidence interval [CI] 95%) exceeded 1 in both datasets, indicating the action of purifying selection. Additionally, for deletions overlapping enhancer sites the OR significantly exceeded 1 in the 1000GP dataset, while the lower CI boundary of the OR was nearly significant, at 0.995, in the ADNI dataset. All intronic and intergenic genomic compartment sets for DHS or enhancer features had mean odds ratios >1 (except ADNI intronic enhancers at 0.96). *SI Appendix*, Table S6c1, (*Note S6c*) lists the odds ratios found in the logistic regressions. These results suggest that purifying selection may be preserving DHS and enhancer epigenomic features by reducing allele frequencies of overlapping deletions. On the other hand, there is a lack of consistent allele frequency shift for genomic compartment sets for transcribed, polycomb-repressed, and heterochromatin features in both deletion datasets, with the mean OR sometimes falling below 1 and the OR CI often extending below 1. In light of the insufficient evidence across datasets for an excess of rare alleles for these features, combined with the lack of reduction in variation described above, we focused the analysis below on DHS and enhancer epigenomic features which showed statistical significance of both key signatures of broad selection against overlapping deletions.

### Differential Selection on Preserving Cellular Activity

The results described above have indicated that purifying selection is acting against the total cellular pleiotropic burden (PlyRS_sum_) of noncoding deletions, preserving both DHS and enhancer regulatory sites. However, these analyses do not clarify if purifying selection preserves DHS or enhancer sites of both tissue/cell-type-specific activity and cellularly pleiotropic activity. One possibility is that deletions removing regulatory elements active in multiple tissues/cell-types incur a greater fitness cost. Another possibility is that since tissue/cell-type-specific elements are vital to organismal development, deletions removing them are subject to a stronger selective effect. It could also be the case that purifying selective pressure on deletions is acting to preserve both types of regulatory sites simultaneously. To distinguish between these scenarios, we calculated two additional PlyRS measures, PlyRS_sum-mono_ and PlyRS_sum-pleio_ (*SI Appendix, Note S1c*). PlyRS_sum-mono_ included the sum of PlyRS values of each deleted base-pair for which a base-pair is only associated with regulatory activity in one tissue/cell-type. PlyRS_sum-pleio_ included the sum of PlyRS values of each deleted base-pair for which that base-pair is associated with regulatory activity in more than one tissue/cell-type. The sum of these two components is the original measure of total cellular pleiotropic burden, PlyRS_sum_. With these additional PlyRS measures, we performed the same analyses as above to examine both a potential reduction in variation and a shift in allele frequency, now applied separately to each component of PlyRS_sum_. This allowed us to determine, within the same sets of real deletions, which scenario of regulatory activity preservation was contributing to the signal of depletion in variation and shift in the AFS as found above.

Fig. 2A shows a significant depletion of variation for DHS or enhancer sites corresponding to both tissue/cell-type-specific activity and for cellularly pleiotropic activity in both 1000GP and ADNI datasets. The effect size of this reduction in variation for PlyRS_sum-mono_ or PlyRS_sum-pleio_ was greater for PlyRS_sum-pleio_ for both noncoding regulatory features, except for enhancer sites in ADNI deletions where the effect size was comparable (error bars overlapping). *SI Appendix*, Tables S5d2-S5d3 (*Note S5d*) lists the effect sizes found in the depletion simulations, including those for intronic and intergenic compartments where depletion values did not consistently favor greater reduction of PlyRS_sum-pleio_. Fig. 2B shows that the magnitude of deletion depletion overlapping DHS or enhancer sites leads to a significantly shifted AFS at both sites of tissue/cell-type-specific activity and cellularly pleiotropic activity. For DHS or enhancer sites in all genomic compartments, the mean odds ratios of the magnitude of depletion for PlyRS_sum-mono_ or PlyRS_sum-pleio_ in association to allele frequency were >1 in both deletion datasets (except ADNI intronic enhancers), and were comparable between PlyRS_sum-mono_ and PlyRS_sum-pleio_. *SI Appendix*, Tables S6c2-S6c3 (*Note S6c*) lists the odds ratios found in the logistic regressions. These results collectively indicate that purifying selection is acting to preserve DHS or enhancer sites of tissue/cell-type-specific activity as well as cellularly pleiotropic activity.

**Fig. 2.**
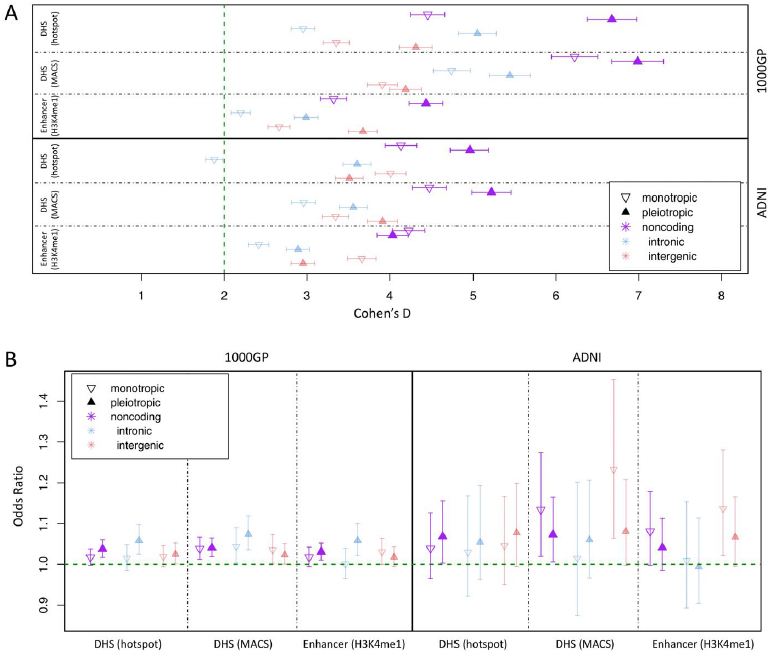
Depletion of deletions and shift of allele frequency spectrum overlapping DHS or enhancer sites of variable cellular activity. (A) We calculated PlyRS_sum-mono_ (monotropic) and PlyRS_sum-pleio_ (pleiotropic) (*SI Appendix, Note S1c*) for every deletion to quantify overlap with DHS or enhancer sites. We plot the degree of reduction in PlyRS_sum-mono_ (or PlyRS_sum-pleio_) for real deletions relative to simulation measured using Cohen’s D (with 95% confidence intervals on the mean reduction). (B) For each deletion we determined the magnitude of PlyRS_sum-mono_ (monotropic) (or PlyRS_sum-pleio_ [pleiotropic]) depletion, calculated as a ratio between its PlyRS_sum-mono_ (or PlyRS_sum-pleio_) and the average PlyRS_sum-mono_ (or PlyRS_sum-pleio_) of its length-matched simulated deletions (*SI Appendix, Note S6b*), for DHS or enhancer sites. We tested whether PlyRS_sum-mono_ (or PlyRS_sum-pleio_) depletion magnitude depends on allele frequency (deletions categorized as rare [AF<=1%] or common), using multivariate logistic regression in the presence of genomic covariates (*SI Appendix, Note S6a*). We plot the regression odds ratio with 95% profile likelihood-based confidence intervals.

### Purifying Selection on CTCF Sites within TAD-loops

We also investigated whether there was evidence of depletion of variation and a shift in the AFS of deletions overlapping topologically associated domain loops (TAD-loops). These large regions of self-interacting DNA facilitate cis-regulatory effects at a wider scale than that of individual regulators (26, 27) and so deletions removing a TAD-loop boundary may be under strong purifying natural selection to preserve the TAD-loop integrity. The distance between TAD-loop boundaries is greater than our longest deletions (25kb limit, [*SI Appendix, Note S2c*]), and consequently deletions in our datasets can only overlap with at most one TAD-loop boundary. Additionally, the TAD-loop boundary data (*SI Appendix, Note S4b*) are less precise than chromatin accessibility or histone modification annotations, so the number of base-pairs of a deletion overlapping a TAD-loop boundary may not reflect actual deleteriousness of the mutation but rather correspond to imprecise annotations on the edges. These characteristics of TAD-loop boundary annotation mean that using PlyRS_sum_ to define the total cellular pleiotropy of overlapping deletions can propagate a potential bias in the measure. To avoid this and still test whether purifying selection may be operating on deletions overlapping TAD-loop boundaries, we measured overlap both as a binary variable and by calculating the maximal PlyRS value (PlyRS_max_, *SI Appendix, Note S1c*) along the length of an overlapping deletion. We performed the same analyses as for the chromatin accessibility or histone modification annotations (*SI Appendix, Notes S5a-S5c*).

Rao Huntley et al. (24) identified that a large majority (86%) of TAD-loop loci had binding from the insulator protein CTCF, which ensures integrity of DNA loops, and consequently, TAD-loop fidelity (28, 29). Given this critical function of CTCF and its presence within most TAD-loop boundaries, we suspected that deletions that overlap TAD-loop loci might be under stronger purifying selection if a deletion also simultaneously overlaps a CTCF site within the TAD-loop boundary (*SI Appendix, Note S4c*). To elucidate this, in addition to identifying the full set of deletions overlapping TAD-loop annotation (TAD-loop), we further refined deletions into two subsets (*SI Appendix, Note S5c*): deletions overlapping TAD-loop but not simultaneously overlapping a CTCF binding site (TAD-loop_noCTCF_) and deletions overlapping TAD-loop while simultaneously overlapping a CTCF binding site (TAD-loop_CTCF_). Only about 1% of all deletions in our datasets overlapped TAD-loop_CTCF_, so we ignored intronic and intergenic designations in the analysis (but maintained them in simulations).

Fig. 3A shows the effect sizes of binary overlap or PlyRS_max_ overlap from comparing real deletions to simulations and indicates that, with respect to the full set of TAD-loops being overlapped (irrespective of whether CTCF sites are simultaneously overlapped), there was minimal depletion of deletion variation, if any. However, as also seen in Fig. 3A, separation into TAD-loop_noCTCF_ and TAD-loop_CTCF_ subsets revealed that a signal of depletion was evident only for deletions overlapping TAD-loop_CTCF_. Deletions in the ADNI dataset exhibited the same characteristic pattern of greater reduction in variation in TAD-loop_CTCF_ versus TAD-loop_noCTCF_ as was seen in the 1000GP dataset; however, the reduction seen in ADNI deletions overlapping TAD-loop_CTCF_ was not statistically significant. We did not find any difference between the effect size of depletion for binary overlap compared to the PlyRS_max_ overlap measure, suggesting that there may not be stronger selection against deletions overlapping the most cellularly pleiotropic TAD-loop_CTCF_. *SI Appendix*, Tables S5d4-S5d5 (*Note S5d*) lists the effect sizes found in the TAD-loop depletion simulations. We also examined whether the depletion magnitude of binary overlap or PlyRS_max_ overlap at TAD-loop loci exhibited dependence on allele frequency using the same logistic regression framework as above with chromatin accessibility and histone modification annotations. Fig. 3B shows compelling evidence of a shift in the deletion AFS based on the magnitude of depletion at TAD-loop_CTCF_, for which the mean odds ratio estimate for binary overlap in 1000GP was 2.70 (minimum [min] 95% CI: 1.35) and in ADNI was 7.67 (min CI: 1.76). The mean odds ratio estimate for PlyRS_max_ overlap of TAD-loop_CTCF_ in 1000GP was 36.80 (min CI: 3.49) and in ADNI was 30.11 (min CI: 1.27). The excess of rare alleles overlapping TAD-loop_CTCF_ dramatically exceeded the shift for TAD-loop_noCTCF_, which displayed only a modest effect in the ADNI dataset (min CI: 1.02) and was not significant in the 1000GP dataset. These results collectively suggest that purifying selection may be acting to preserve TAD-loop integrity by specifically preserving CTCF binding motifs within TAD-loop boundaries. *SI Appendix*, Tables S6c4-S6c5 (*Note S6c*) lists the odds ratios found in the TAD-loop logistic regressions.

**Fig. 3.**
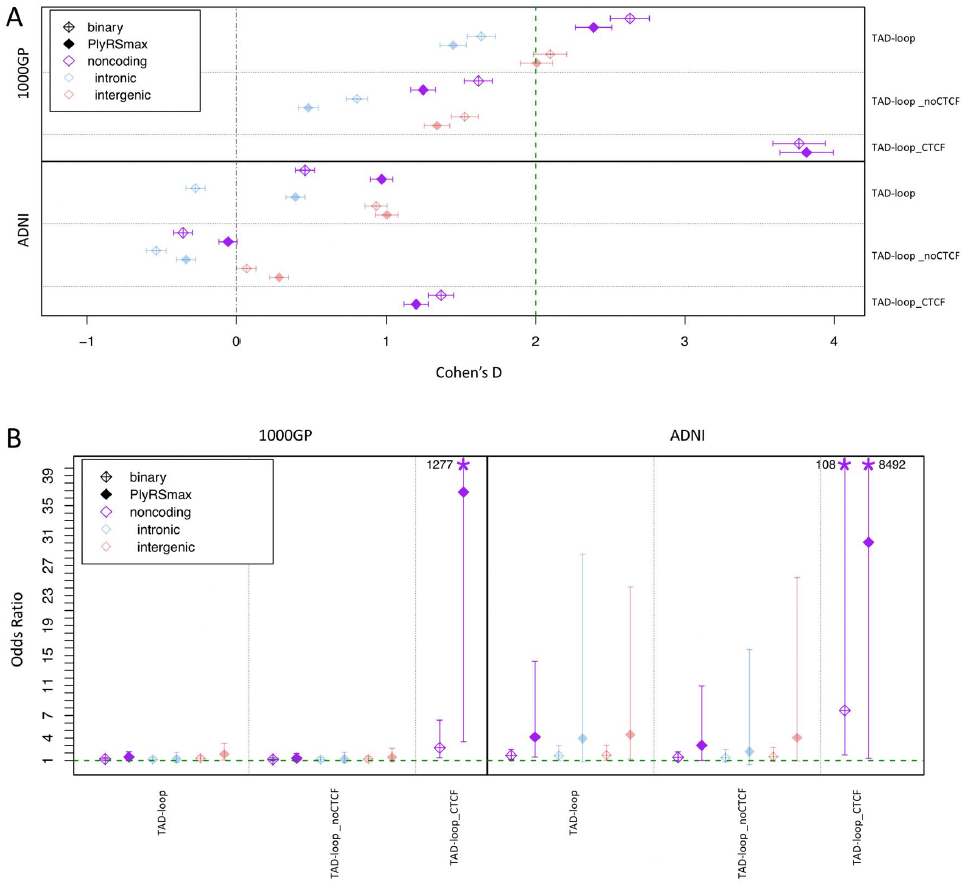
Depletion of deletions and shift of allele frequency spectrum overlapping TAD-loop regulatory sites. (A) We calculated a binary variable and PlyRS_max_ (*SI Appendix, Note S1c*) for every deletion to quantify overlap with sites of TAD-loop. We plot the degree of reduction in the binary variable (or PlyRS_max_) for real deletions relative to simulation measured using Cohen’s D (with 95% confidence intervals on the mean reduction). (B) For each deletion we determined the magnitude of binary variable (or PlyRS_max_) depletion, calculated as the difference between the binary variable (or PlyRS_max_) and the average binary variable (or PlyRS_max_) of its length-matched simulated deletions (*SI Appendix, Note S6b*), for sites of TAD-loop. We tested whether binary variable (or PlyRS_max_) depletion magnitude depends on allele frequency (deletions categorized as rare [AF<=1%] or common), using multivariate logistic regression in the presence of genomic covariates (*SI Appendix, Note S6a*). We plot the regression odds ratio (OR) with 95% profile likelihood-based confidence intervals.

## Discussion

Using the clarity of genomic deletions to identify loss of noncoding regulatory function, we have examined whether purifying selection is operating to preserve noncoding regulatory sites of chromatin accessibility (DHS), histone modification (enhancer, transcribed, polycomb-repressed, and heterochromatin), and topologically associated domain loops (TAD-loops). Analysis of selection in the noncoding genome is motivated by prior findings in human genetics from genome-wide association studies that conclude most of heritability is due to relatively common noncoding alleles within regulatory annotations (3). Initially, these findings appeared inconsistent with the expectation that disease-associated alleles are under pressure from purifying selection. However, recent studies demonstrated that complex trait effect sizes are negatively correlated with allele frequency, hinting at the action of purifying selection (30–32). These observations put the question of the effect of noncoding regulatory alleles on function and fitness at the forefront of genomic studies ranging from basic evolutionary genetics to the allelic architecture of common human traits. Since a principal characteristic of human regulatory element function is their non-uniform activity across tissues and cell types, interpreting fitness consequences from genetic variants in noncoding regions is thus inherently linked to corresponding regulatory element cellular activity. To incorporate this defining feature into the study of noncoding purifying selection, we have developed a statistical method, Pleiotropy Ratio Score (PlyRS), which quantifies the extent of abundance of cellularly pleiotropic activity for individual base-pairs.

Using our PlyRS method, our results indicate that purifying selection acts on both DHS and enhancer sites, as evident by both the depletion of deletions overlapping these annotations and a shift in the allele frequency spectrum of overlapping deletions towards rare alleles. Using simulated deletions in a length-matched framework and covariate-aware analyses, we notably found statistically significant evidence at DHS or enhancer regions that both sites of tissue/cell-type-specific activity and sites of cellularly pleiotropic activity are preserved by selection. We find some evidence that cellularly pleiotropic variants may be subject to a stronger reduction in variation than cell-type-specific variants. However, ambiguity between tissue/cell-type-specific and cellularly pleiotropic sites in terms of AFS shifts indicates that the strength of purifying selection across both types of regulatory site cellular activities may be roughly equivalent. Additional analysis on larger datasets would be needed to accurately quantify the relative contributions of selection on sites of variable regulatory activity.

In contrast to the findings above, we did not find evidence of purifying selection acting on other epigenomic annotations such as transcribed, polycomb-repressed, or heterochromatin sites, consistent with previously reported findings (14, 16). In the absence of statistical confirmation, we can conclude that, notwithstanding any specific regulatory locus potentially being under selective constraint, these classes of epigenomic regulators as a whole are not selectively preserved in noncoding space. These results underscore the importance of DHS and enhancer annotations for specifying critical cellular regulation. Notably, our findings parallel the observation in human genetics that the largest fraction of heritability resides in regulatory space marked by DHS or enhancer features (3, 33). Our results additionally support the hypothesis that an aggregate selective burden may occur on long deletions that overlap multiple DHS or enhancer sites simultaneously (14, 34). We find suggestive evidence that this may be the case for deletions longer than the median length in our datasets, especially on those that overlap cellularly pleiotropic sites (*SI Appendix*, Fig. S4).

We have also presented evidence that purifying selection is operating to preserve TAD-loop boundary integrity by preserving co-localized CTCF binding sites. However, we did not find statistical evidence that selection is acting against deletions overlapping TAD-loop boundaries without simultaneous removal of CTCF sites. We found conclusive statistically significant evidence for this preservation of TAD-loop_CTCF_ sites in 1000GP but only a qualitative trend for this in ADNI. The difference in significance for these findings between deletion datasets may simply be due to the difference in power to see this effect, as there are 4x the number of deletions in 1000GP in comparison to ADNI. We did not find statistically significant evidence in either dataset that the sites of highest cellular pleiotropy of TAD-loop_CTCF_ provides additional signal for purifying selection beyond that for TAD-loop_CTCF_ sites of any cellular pleiotropy. This equivalence may again be due to lack of power: either five primary tissues/cell-types in TAD-loop boundary analysis are not numerous enough to see a difference (compared to the 25 primary tissues/cell-types used in the analysis of chromatin accessibility and histone modification features), or deletions overlapping cellularly pleiotropic TAD-loops are already so few in number that power is limited (only 4% [1000GP] or 8% [ADNI] of all deletions in our datasets). As with the DHS and enhancer findings mentioned above, larger datasets may provide the power needed to clarify the relative contributions of selection on TAD-loop and CTCF sites of variable activity, as well as provide better resolution of TAD compartments versus TAD-loop boundaries which may improve analyses. The PlyRS method is flexible and easily allows for the addition of new and larger regulatory datasets as they become available.

## Materials and Methods

We used deletions from two datasets, the 1000 Genomes Project (1000GP, [(14)] and the Alzheimer’s Disease Neuroimaging Initiative (ADNI [(22)], *SI Appendix, Note S3*), to examine selective constraint within regulatory regions. The two deletion datasets have different callset properties, enabling robustness of the analysis. 1000GP consists of deletions derived from low-coverage whole genome sequencing (WGS) that span a wider length range and are genotyped from individuals of diverse demographic histories. ADNI consists of deletions derived from high-coverage WGS data that are on average longer and more rare, using genotypes from the subset of individuals that we determined were of European ancestry as identified by principal components analysis. For both deletion datasets, we restricted our analyses to noncoding deletions by removing any deletion that overlapped any exon or UTR by one base-pair or more, as exonic deletions have been previously shown to be under strong purifying selection because of their protein-altering effects (35). We also examined only deletions occurring on autosomes because sex-chromosome functional elements may involve complex sex-biased regulation (36) which might be subject to unique selective properties. To mitigate non-uniform (i.e. biased) deletion callability in the noncoding genome which might distort the AFS of the remaining set of deletions, we additionally excluded deletions overlapping any regions of low mappability, segmental duplications, centromeres, and reference assembly gaps. Additional details on the deletion datasets and filtering criteria are given in *SI Appendix, Note S2*. Specific characteristics of the deletion datasets are shown in *SI Appendix*, Table S1 (1000GP) and Table S2 (ADNI). An extended description of the ADNI dataset construction process is given in *SI Appendix, Note S3*. Information on obtaining ADNI data access, including files we deposited [in-progress] for this project, can be found at: http://adni.loni.usc.edu/data-samples/access-data/.

We used regulatory data from the NIH Roadmap Epigenomics Consortium (REC) for definition of regulatory breakpoints as well as uniform processing across multiple tissue/cell-types (2). We use annotation data for sites of chromatin accessibility (DNase I hypersensitivity “DHS”) and histone modification (H3K4me1 “enhancer”, H3K36me3 “transcribed”, H3K27me3 “polycomb-repressed”, and H3K9me3 “heterochromatin”). Two sets of DHS annotation (hotspot and MACS) were used to check for consistency. We used all 25 primary tissues/cell-types (*SI Appendix, Note S4a* and Table S3) for which data were available across all six callsets for each tissue/cell-type. We additionally used TAD-loop boundary regulatory data consisting of a callset of 5 primary tissues/cell-types ([(24)], *SI Appendix, Note S4b*). Additional details on the regulatory datasets are given in *SI Appendix, Note S4*. Identity of the tissues and cell-types analyzed from REC is shown in *SI Appendix*, Table S3. For all analyses involving DNase hypersensitivity or histone modification regulatory features, we excluded deletions (and genomic space) overlapping TAD-loop boundaries (*SI Appendix, Note S5c*), as deletions disrupting TAD-loop integrity may already be under purifying selection owing to the potentially resulting cis-regulatory effects. In this way, we ensure reliable interpretation of selective effects on deletions disrupting chromatin accessibility or histone modification, without introducing potential confounding from selective pressure from co-localized TAD-loop disruption, which we analyzed separately.

To examine potential purifying selection against deletions to preserve regulatory features, we examined deletion overlap in the context of regulatory tissue activity. To properly “count” tissue activity removed by deletions overlapping regulatory features, we developed a statistical method called Pleiotropy Ratio Score (PlyRS), which calculates a correlation-adjusted count of cellular pleiotropy for each base-pair in the noncoding genome. A description of the calculation of PlyRS, and derived PlyRS measures calculated for deletions, is given in *SI Appendix, Note S1*. Source code of PlyRS calculation can be downloaded from the repository [in-progress] on Github: https://github.com/davidwradke/PlyRS.

To determine if the action of purifying selection is occurring against deletions overlapping regulatory sites, we required the identification of two key signatures: reduction of genetic variation overlapping the sites and a shift in the allele frequency spectrum (AFS) towards rare variants of the remaining alleles overlapping the sites. These signatures were assessed in light of results from deletion simulations. A description of the simulation procedure and significance calculation of reduction in variation is given in *SI Appendix, Note S5*. Descriptions of the procedure involving multivariate regression on deletion genomic covariates and significance calculation of shift in allele frequency spectrum are given in *SI Appendix, Note S6*.

## ACKNOWLEDGMENTS

We thank members of the Sunyaev lab and Matt Maurano for helpful discussion on the topic. We thank Sheila Sutti Diamond for administrative help related to the Alzheimer’s Disease Neuroimaging Initiative data used and generated in this project. We also thank Sung Chun, Nikolaos Patsopoulos, Deb Farlow, and Ruth McCole for helpful discussion related to the ADNI work. We thank the ADNI individuals for their participation in the consortium.

This material is based upon work supported by the National Science Foundation Graduate Research Fellowship Program under Grant No. DGE1144152 (D.W.R.). Any opinions, findings, and conclusions or recommendations expressed in this material are those of the author(s) and do not necessarily reflect the views of the National Science Foundation. D.W.R. was also supported by the National Institutes of Health (NIH) under Ruth L. Kirschstein National Research Service Awards 4T32HG002295-14 and 5T15LM007092-27 and is currently supported by NIH Grant 5R01HG010372. S.R.S. is currently supported by NIH Grants R35GM127131, R01MH101244 and U01HG006500. R.C.G is currently supported by NIH Grants R01-HG009922 and R01-HL143295 and by the Franca Sozzanni Fund.

Data collection and sharing for this project was funded by the Alzheimer’s Disease Neuroimaging Initiative (ADNI) (National Institutes of Health Grant U01 AG024904) and DOD ADNI (Department of Defense award number W81XWH-12-2-0012). ADNI is funded by the National Institute on Aging, the National Institute of Biomedical Imaging and Bioengineering, and through generous contributions from the following: AbbVie, Alzheimer’s Association; Alzheimer’s Drug Discovery Foundation; Araclon Biotech; BioClinica, Inc.; Biogen; Bristol-Myers Squibb Company; CereSpir, Inc.; Cogstate; Eisai Inc.; Elan Pharmaceuticals, Inc.; Eli Lilly and Company; EuroImmun; F. Hoffmann-La Roche Ltd and its affiliated company Genentech, Inc.; Fujirebio; GE Healthcare; IXICO Ltd.; Janssen Alzheimer Immunotherapy Research Development, LLC.; Johnson Johnson Pharmaceutical Research Development LLC.; Lumosity; Lundbeck; Merck Co., Inc.; Meso Scale Diagnostics, LLC.; NeuroRx Research; Neurotrack Technologies; Novartis Pharmaceuticals Corporation; Pfizer Inc.; Piramal Imaging; Servier; Takeda Pharmaceutical Company; and Transition Therapeutics.

The Canadian Institutes of Health Research is providing funds to support ADNI clinical sites in Canada. Private sector contributions are facilitated by the Foundation for the National Institutes of Health (www.fnih.org). The grantee organization is the Northern California Institute for Research and Education, and the study is coordinated by the Alzheimer’s Therapeutic Research Institute at the University of Southern California. ADNI data are disseminated by the Laboratory for Neuro Imaging at the University of Southern California.

