## Supplementary Information for "Purifying selection on noncoding deletions of human regulatory elements detected using their cellular pleiotropy"

[a]Program in Genetics and Genomics, Biological and Biomedical Sciences PhD Program, Harvard Medical School, Boston, MA, 02115, USA; [b]Department of Biomedical Informatics, Harvard Medical School, Boston, MA, 02115, USA; [c]Division of Genetics, Department of Medicine, Brigham and Women's Hospital and Harvard Medical School, Boston, MA, 02115, USA; [d]Broad Institute of Harvard and MIT, Cambridge, MA, 02142 USA; [e]Department of Psychiatry and Biobehavioral Sciences, University of California, Los Angeles, CA, 90095, USA; [f]Department of Organismic and Evolutionary Biology, Harvard University, Cambridge MA, 02138, USA; *Data used in preparation of this article were obtained from the Alzheimer's Disease Neuroimaging Initiative (ADNI) database (adni.loni.usc.edu). As such, the investigators within the ADNI contributed to the design and implementation of ADNI and/or provided data but did not participate in analysis or writing of this report. A complete listing of ADNI investigators can be found at: http://adni.loni.usc.edu/wp-content/uploads/how_to_apply/ADNI_Acknowledgement_List.pdf

**This PDF file includes:**

Supplementary text
Figures S1 to S4
Tables S1 to S3
SI References

#### Note S1

**Measurement of deletion horizontal and vertical 'axes'**

##### Note S1a

***Horizontal axis: base-pairs deleted***

To examine how noncoding deletions can potentially remove regulatory function at a genomic locus along the horizontal axis, we calculate a count of the number of regulatory elements overlapped by a deletion. A single deletion, depending on its length, may remove one, or more adjacent, regulatory elements. There is a mismatch, however, between start and end coordinates of most pleiotropic regulatory elements when compared across tissues and cell-types because the regulatory element annotation may vary slightly in base-pairs (bp) length. Regulatory annotations would ideally have the same number of contiguous nucleotides identified at each locus across active tissues and cell-types. However, this is not the case with real data due to the ChIP-seq experimental signal not providing precise 'peak' calling of the regulatory elements (SI 1). Therefore, since we cannot confidently determine which coordinates are correct or incorrect (as to the 'peak') in any one tissue versus another, we use the specific bp annotation from narrow regions of enrichment ($p<=0.01$) for histone modification data and DNase I hypersensitivity data (MACS peak caller [SI 2], NarrowPeak) and additionally the specific bp annotation from general-sized regions of DNA accessibility ($p<=0.01$) for DNase I hypersensitivity data (hotspot algorithm [SI 3], BroadPeak) (located at: Roadmap supplement website). The bp annotation therefore provides a 'cloud' over the real regulatory element co-localization and does not require fixing precise coordinate boundaries between each tissue/cell-type. Because the annotation is at the per-bp level, regulatory element annotations can be equally compared across tissues/cell-types. When examining the horizontal axis, we therefore use the number of regulator-annotated bp deleted found within at least one tissue ("bp_affected"), only examining this on a per-annotation basis. While bp_affected does not correspond to the number of regulatory elements deleted, it instead corresponds to the approximate scaled fraction of regulatory elements deleted. For example, if bp_affected = 15, this might correspond to roughly 1/10 of the regulatory element being deleted (using an average regulatory element coordinate length of 150bp [SI 3]). Since even partial deletion of a regulatory element might render it at least partially inoperable due to altered binding, bp_affected therefore serves as a scaled proxy of real regulatory element removal.

##### Note S1b

***Vertical axis: PlyRS calculation***

To examine how noncoding deletions can potentially remove regulatory function at a genomic locus along the vertical axis, we calculate a correlation-adjusted count of regulatory element activity amongst the tissues/cell-types analyzed. A simple count of regulatory activity would be highly influenced by the input tissues/cell-types, and may under/over-represent truly pleiotropic regulatory activity across cell-types if a subset of highly correlated tissues/cell-types (such as blood cells) were to dominate the dataset. For a hierarchical clustering of tissues example using enhancer sites, see Kundaje et al. Fig. 6A (SI 1). Instead of a simple count, we calculate an activity correlation-adjusted count of regulatory element activity across tissues/cell-types, and scale this from 0 (representing no regulatory activity in any tissue/cell-type) to 1 (representing regulatory activity in all tissues/cell-types). This count is calculated per-base-pair from the regulatory annotations used (SI Note S4). We call this count the Pleiotropy Ratio Score ("PlyRS", pronounced 'ply-ers' as in the pliers hand tool).

For derivation of PlyRS, we adapted the PSIC method (SI 4) which was originally developed as a method to assess 'independent counts' when looking at a multiple sequence alignment of amino acid substitutions. PSIC is a major component of the widely used PolyPhen-2 tool (SI 5). Fig. S1b1 provides a helpful schematic of the derivation of PlyRS.

**Figure S1b1: Schematic of derivation of PlyRS.**

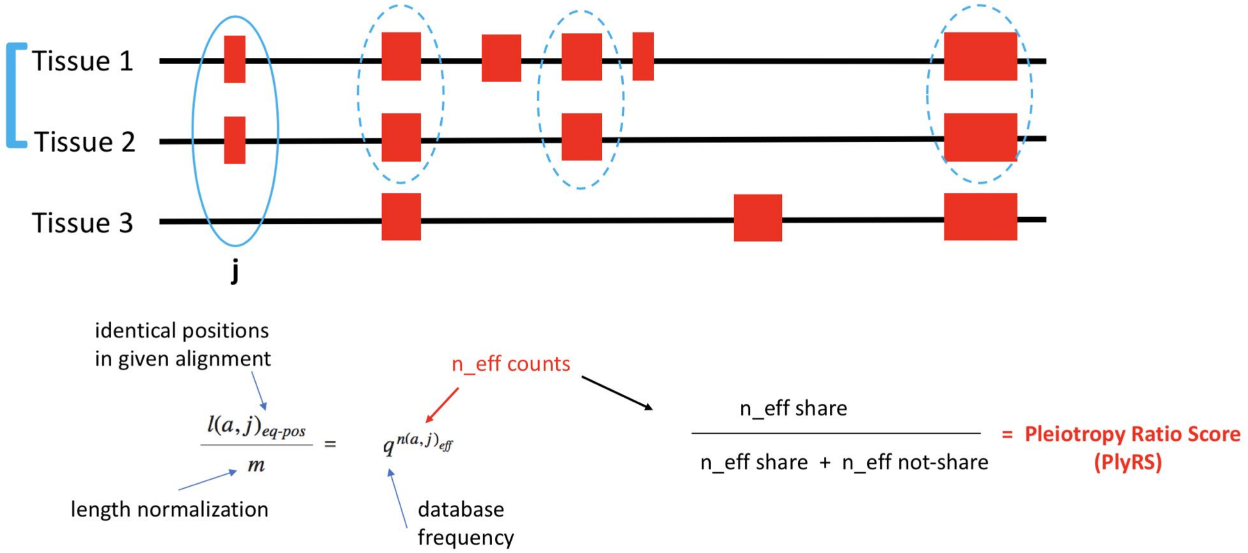


For a set of tissues analyzed (here, three) we find the tissues that share a regulatory element annotation at a particular base-pair (here, position j has Tissue 1 and Tissue 2 sharing a regulatory annotation). We count the number of identical sharing positions throughout the rest of the genome (in our case, only autosomal positions are analyzed [SI Note S2c]). From these values we can calculate the fraction on the left side of the equation. The number of identical sharing positions forms the numerator, and the number of positions analyzed genome-wide forms the denominator. From this fraction, we use a relation to a Bernoulli experiment. The central idea is to equate the frequency of identical positions in a real alignment with the probability of identical alignment given random Bernoulli sequences, by solving for n_effective counts. The database frequency (here, q) is the frequency of the regulatory annotation measured across all tissues and sites genome-wide. We can then solve for n_eff counts, which gives us an adjusted count based on the underlying correlation between the tissues analyzed. This number in principle will not be very informative as a raw number, since it would scale very differently at each base-pair depending on the set of tissues sharing a regulatory element annotation at each coordinate, especially when the number of tissues analyzed is large (in our case, 25). Therefore, we repeat this process for all the tissues that originally did not share a regulatory annotation at a particular position (here position j has Tissue 3 not sharing a regulatory annotation). We calculate n_eff counts for this set of tissues. From these two n_eff counts ('n_eff share' and 'n_eff not-share') we can derive the Pleiotropy Ratio Score (PlyRS) by taking the n_eff counts from sharing and divide by the total n_eff counts from both sharing and not sharing. SI Fig. S2 displays example PlyRS compared to tissue/cell-type raw counts. SI Fig. S3 displays example PlyRS values (related back to an otherwise standard count of 1) when regulators are annotated as tissue-specific or cell-type-specific.

For our purposes, we define a deletion at a regulatory element as having simultaneously both a horizontal axis component and vertical axis component, even if that deletion overlaps a regulator with only single tissue/cell-type-specific activity (in the vertical axis). This definition is needed for the vertical axis since we are only able to observe potential pleiotropy from the tissues/cell-types analyzed in our dataset, however, it may be the case that had an additional tissue(s) been analyzed, pleiotropy would have been found at a regulatory locus that otherwise appears as tissue/cell-type-specific in our current dataset. Therefore, in our definition, the bottom count of the vertical axis would be from tissue/cell-type-specific regulators (using its PlyRS), since we can't exclude that those loci may actually have additional cellular pleiotropy currently hidden given the incomplete set of bodily tissues currently analyzed. Therefore, while a certain regulatory locus may be tissue/cell-type-specific (i.e. not cellularly pleiotropic in definitional sense), it will nonetheless have a Pleiotropy Ratio Score (PlyRS) value > 0. A PlyRS value = 0 corresponds to base-pairs for which there is no annotated regulator at that genomic position in any tissue/cell-type analyzed in the total set of tissues/cell-types.

#### Note S1c

***Pleiotropy Ratio Score (PlyRS) measures***

***PlyRS$_{sum}$:*** corresponds to the total cellular pleiotropy (for a specific regulatory feature) of the deletion, encompassing both the horizontal and vertical 'axes' along which purifying selection may be operating on the deletion. It is calculated by summing together all PlyRS values found along the length of the deletion (this is the same result as multiplying the number of overlapped regulatory base-pairs by the average PlyRS value found along the deletion). PlyRS$_{sum\text{-}mono}$ and PlyRS$_{sum\text{-}pleio}$ component sums combined equals the measure of total cellular pleiotropic burden. This value correlates strongly with the total number of base-pairs (and also with the number of regulatory base-pairs) of the deletion, since the horizontal axis often forms the bulk of the sum because tissue/cell-type-specific sites make up roughly a quarter of all regulatory sites.

**PlyRS$_{sum\text{-}mono}$:** includes the sum of PlyRS values of each deleted base-pair for which that base-pair is only associated with regulatory activity in one tissue/cell-type. The count at this base-pair is not 1, however, because the count is adjusted by the correlation between all the tissues and cell-types being analyzed (SI Note S1b).

**PlyRS$_{sum\text{-}pleio}$:** includes the sum of PlyRS values of each deleted base-pair for which that base-pair is associated with regulatory activity in more than one tissue/cell-type.

**PlyRS$_{max}$:** corresponds to the maximal PlyRS value found at any base-pair from examining all base-pairs along the length of a deletion. This value can a maximum of 1, representing 100%

cellular pleiotropy across the tissues and cell-types analyzed. This measure is more stable (than $PlyRS_{sum}$) with regulatory annotations that have less precision on boundaries (such as TAD-loops).

#### Note S2

**Deletion datasets**

##### Note S2a

***1000 Genomes Project phase 3 (1000GP)***

The 1000 Genomes Project Consortium phase 3 (1000GP) SV callset (SI 6) was downloaded from the FTP site hosted by EBI (ftp://ftp.1000genomes.ebi.ac.uk/vol1/ftp/). This dataset was derived from running multiple structural variant algorithms on low-coverage (~7x average coverage) whole genome sequencing data of 2,504 individuals from 26 populations. The VCF file containing the deletion calls with allele frequency (AF) was located in subdirectory '/phase3/integrated_sv_map/ALL.wgs.mergedSV.v8.20130502.svs.genotypes.vcf.gz' ('original file') and did not include any Y chromosome calls. Deletion calls passing metrics for fine-resolution of breakpoints are contained in TABLE_3-Breakpoints.xlsx and were additionally downloaded as a text file from subdirectory 'phase3/integrated_sv_map/supporting/breakpoints/1000GP_phase3_all_bkpts.v5.txt.gz' ('breakpoints file'). These data were generated from a variety of published deletion callers and underwent quality control for accuracy in variant genotyping and fine-resolution of coordinate breakpoints. Since we want to analyze only variant calls which are deletions (i.e. loss) of genomic information relative to the human reference genome, not analyze the absence of an insertion of genomic information (potentially human reference sequence insertions in the individuals catalogued), only variant calls likely to be true deletions from BreakSeq (SI 7) prediction were retained (only "NAHR", "NAHR_EXT", or "NH" MUTMECH designations as specified in the 'breakpoints file'). To arrive at this set with corresponding allele frequency, we extracted AF from the 'original file' containing all the deletion calls and matched deletion name identifiers from this file with that in the 'breakpoints file'. For deletions with the same/synonymous name identifier that had more than one set of breakpoints identified (about 0.2% of deletions), we used the average breakpoint-resolved start coordinate and average breakpoint-resolved end coordinate. The total collection of autosomal deletions with AF gathered in these processes numbered 22,684.

Nearly all 'SVTYPE=DEL_*\t') variants were removed because of this procedure (or from mappability filters-see SI Note S2c), as expected, since these variants were identified by 1000GP as likely human reference sequence mobile element insertions. Remaining 'SVTYPE=DEL_*\t' deletions were not removed, as manually removing these variants may lead to unknown biases in downstream analysis when those same underlying genomic coordinates would otherwise be allowed in regulatory assays or computational simulations (i.e. removing these deletions manually may induce artificial depletion in these loci). We are interested in the missing sequence in an individual of any uniquely-mappable sequence in the noncoding human genome, given our other filters. Since some MEIs (which become called as deletions in some individuals) may have epigenomic regulatory annotation, those 'deletion' variants should be left in the dataset because the absence of that sequence in one or more individuals may have functional consequence in those individuals. We don't judge *a priori* the importance of that noncoding sequence space if it is uniquely alignable and is already available to regulatory experimental assays. For example, if sequence at a locus is marked as having DHS activity, then if a MEI 'deletion' occurs at that locus, that means some humans don't have that open chromatin, which may be functionally important. Ideally, perfectly identified human ancestral sequence would be able to identify true losses of genetic information (derived deletions after last common ancestor) from contemporary deletion data, however reconstructing the human ancestral genome as well as comparison to primate genomes is difficult due to differing reference sequence qualities (SI 8).

The allele frequency used was the global AF ("AF=" in the 'original file'). Most deletions are population specific because they are rare (singleton or doubleton, *etc.*), but common deletions are subject to genetic drift and by taking a global AF, this smooths-out these population-specific demographic histories for those variants. We are interested in studying purifying selection to preserve regulatory elements broadly during human evolution, not examine purifying selection on regulatory elements for potential population-specific deletions. Also, besides losing statistical power by breaking the deletion set into population-specific sets, since each demographic population would have a different genetic distance to the human reference sequence (given the underlying construction of the reference sequence being biased toward European variation (SI 9), differences seen in the underlying population-specific deletion sets would be potentially artifactual and likely not representative of biological differences. Also, with population-specific deletion datasets (and resulting population-specific simulations), we would be re-examining common deletions multiple times (since they would often be shared between most/all populations), which may introduce statistical confounding in the interpretation of the results.

##### Note S2b

***Alzheimer's Disease Neuroimaging Initiative (ADNI)***

We generated a quality-controlled and filtered deletion dataset numbering 10,619 autosomal deletions by running the CNV algorithm GenomeSTRiP (SI 10) on high-coverage (~42x average coverage) whole genome sequencing data of 808 participants in the Alzheimer's Disease Neuroimaging Initiative (ADNI) (SI 11). Deletion calls (as compared to the human reference sequence GRCh37) were retained from individuals of European ancestry (752 individuals) identified from conducting principal components analysis on common SNVs. A variety of quality-control criteria and filters designed to balance deletion counts while also ensuring robust breakpoint accuracy and genotyping accuracy were applied to arrive at a final deletion callset. A full description of our procedures are given in SI Note S3. The deletion dataset with population AF can be downloaded from (link given here when available). It should be noted that though many individuals in the dataset have a phenotype of mild cognitive impairment (342/752) or Alzheimer's Disease (177/752), natural selection would be expected to be mild or nonexistent on variants associated with these phenotypes (unless pleiotropic), as they are post-reproductive in nature (all individuals in the dataset were at least age 50). Nearly all identified deletion variants in the dataset would be unrelated to phenotype labels; however, if any are included, they may actually be viewed as conservative to results in our downstream analyses, since a population cohort with a few disease-associated regulatory deletions would actually slightly bias our depletion and allele frequency spectrum (AFS) shift results in a direction against our conclusion. Therefore, any significant result in our analysis that remains with the inclusion of a few phenotypically-associated variants (if known) would likely be slightly more significant had these deletions been removed from our analysis. We did not find any statistically significant phenotypically-associated deletion variants in ADNI (see SI Note S3e, section: Flat quantile-quantile plot).

#### Note S2c

***Filters applied to both 1000GP and ADNI datasets***

Additional filters were applied to ensure careful examination of the effects of selection on deletions overlapping regulatory elements. We removed deletions overlapping exons. Genomic coordinates used to identify exonic and genic sites were downloaded from Ensembl Biomart (http://grch37.ensembl.org/biomart/martview/ with dataset: 'Human genes (GRCh37.p13)'). We removed all sex-chromosome deletion variants, thereby removing 1000GP X chromosome deletions (there were no X chromosome deletions called in ADNI and no Y chromosome deletions called in either 1000GP or ADNI). BEDTools software (SI 12) version 2.26.0 was used to remove deletion variants that overlapped loci excluded from analysis. Using tracks downloaded from the UCSC Genome Table Browser (https://genome.ucsc.edu/cgi-bin/hgTables), we removed deletions overlapping any regions of low mappability (wgEncodeCrgMapabilityAlign100mer) (see SI Note S2d below), segmental duplications (genomicSuperDups), centromeres and reference assembly gaps (gap). We additionally removed ADNI deletions overlapping regions of B-cell instability using the same regions already excluded by the 1000GP consortium in their released deletion callset (SI 6). Additionally, deletions longer than 25kb were removed because downstream analysis with simulated deletions/mock datasets require independent coordinate assessment and deletions longer than this may cause spurious 're-mutation' in the mock datasets (because of simulation procedure rules used) not representative of deletions in the real datasets. Because of the stringent filters already employed, principally the low mappability filter, only three 1000GP deletions and one ADNI deletion were removed because of this length cutoff.

The resulting deletion datasets remaining after the filtering procedures were applied (including as described below in SI Note S2d) included 12,013 1000GP deletions and 3,306 ADNI deletions. These deletion sequences are found with respect to the human hg19/b37 reference genome. Deletion dataset characteristics are summarized in SI Table S1 (1000GP) and SI Table S2 (ADNI). Allele frequency was kept as the raw deletion AF with respect to the reference, not minor AF (MAF), because we want to analyze the loss of genetic information in the form of deletions, not just the minor allele which might represent the non-deleted state for extremely common deletions, potentially resulting in biases in AF comparison analyses.

Note S2d

***Deletion callability and need for unique coordinates***

Our analysis depends on accurate deletion genotyping and accurate regulatory element annotation. In regions of the genome where sequence is non-unique, (i.e. for any 100bp stretch, that sequence appears in more than one location in the reference human genome), sequence read coverage may be missing, averaged across all sites in the genome, or over-represented depending on alignment algorithm parameters used. This can present problems for deletion calling as well as regulatory element peak calling. In addition, if these sequences are, on-average, less functional (due to repeat sequences), then selection may be operating in a relaxed manner in these regions, biasing or confounding analyses in identifying a shift in the deletion AFS. Additionally, if simulations are performed where deletions are randomly placed along the genome, if non-unique regions are available as potential random placement locations for deletions, a functional-to-less functional 'spreading' will occur. See Fig. S2d1 for an example of 1000GP noncoding deletion spreading in simulations from real (more-unique) coordinates to mock (less-unique) coordinates (using RepeatMasker [SI 13] annotations). This is because in the real deletion callset, non-unique genomic regions presented less-confident deletion evidence, on average, than unique regions of the genome, and are therefore likely underrepresented in the deletion callset. Therefore, more deletions calls were made in unique regions compared to non-unique (all other things being equal) and so in a simulation framework where deletions are randomly placed throughout the genome, there will be a 'migration' from unique-to-non-unique, at a modest noticeable extent. This effect is very hard to control for in matched simulations, given the covariance of non-unique regions with GC content, recombination rate, and other genomic features. This migration of deletion calls can bias analyses that depend on overlap with regulatory annotations (that would predominantly be located in more unique regions of the genome given the same issues encountered for ChIP-seq assay sequence read mapping). Therefore, we decided to ensure robust genomic coordinates for our analyses, and restricted to only unique sites in the genome (i.e. for any 100bp stretch, that sequence appears only in that one location in the human genome).

**Figure S2d1: Deletion 'spreading' in simulations.**

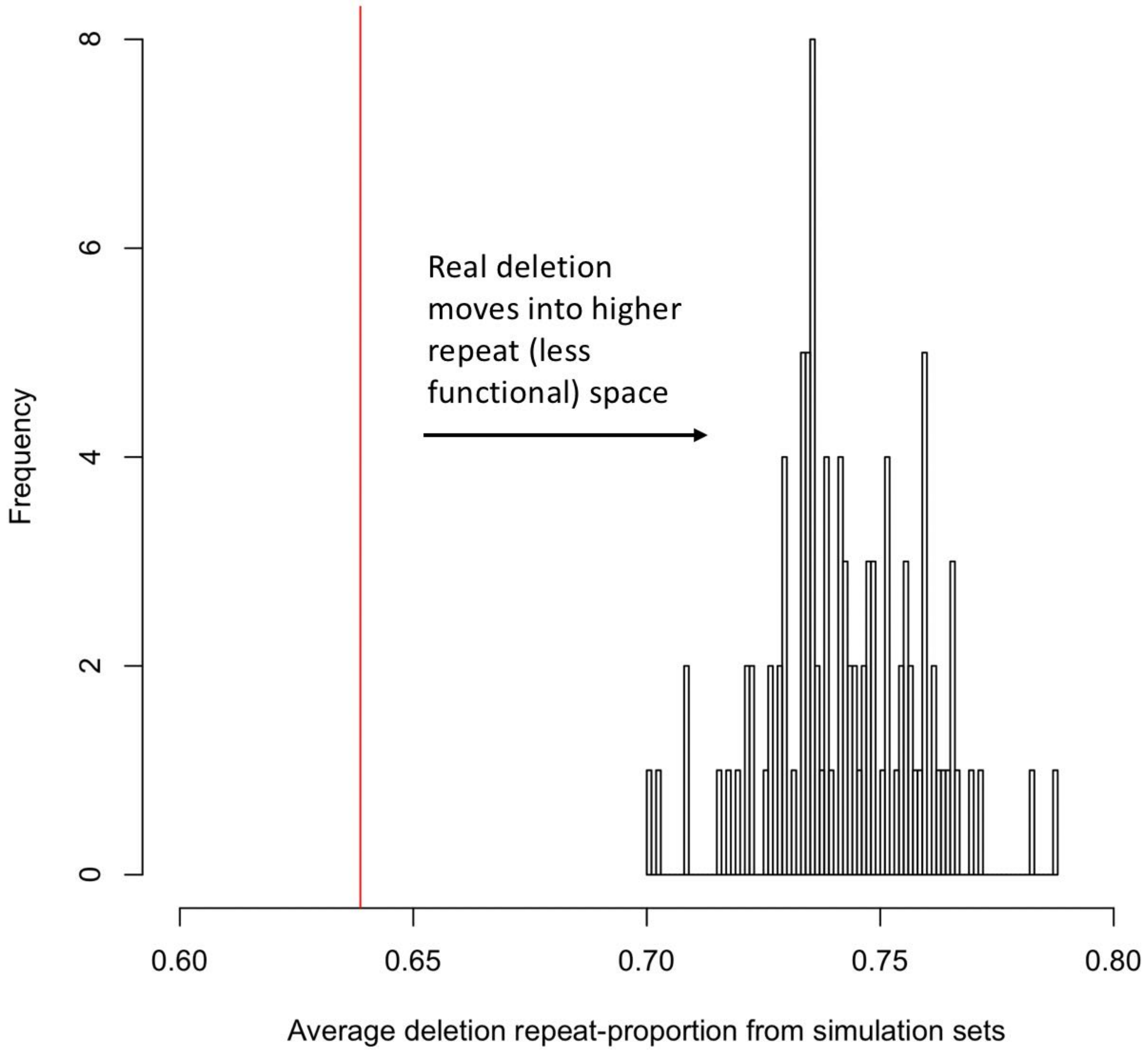


#### Note S3

**ADNI dataset construction**

##### Note S3a

***Brief Summary***

Whole genome sequencing (WGS) was previously performed on >800 participants in the Alzheimer's Disease Neuroimaging Initiative (ADNI). For this project, we compiled a deletion callset on 752 ADNI individuals of European ancestry using the published deletion algorithm GenomeSTRiP (SI 10). We examined re-aligned analysis-ready BAM files (~150 terabytes), which were generated by the Broad Institute using their GATK 'best-practices' pipeline. By combining a variety of computational quality-control criteria and filters on the deletion variants (see SI Note S3c) identified by GenomeSTRiP, we arrived at a callset of 10,619 autosomal deletions (for deletion characteristics see Table S3g2 [SI Note S3g]). A schematic overview of our deletion callset generation is diagrammed in Fig. S3g1 (SI Note S3g). We identified that our deletions calls a) have the qualitatively expected allele frequency distribution (see SI Note S3e, section: Allele frequency spectrum shape) of a robust population dataset, b) have high concordance with 1000 Genomes Project-identified (SI 6) common deletions (>=5%) for deletions at or longer than our median length of 3,114 base-pairs, and c) have a well-behaved, flat quantile-quantile plot distribution, as would be expected from a dataset such as the ADNI WGS which is likely underpowered to identify any single variants of large phenotypic effect, indicating high-quality genotyping across subsets of the ADNI data. We provide three usage cases for which these data may be useful to researchers (see SI Note S3f), and make these data publicly available to registered users of the ADNI (see SI Note S3d).

#### Note S3b

***Background***

It is estimated that by 2050, the prevalence of Alzheimer's Disease (AD), the most common form of mental deterioration amongst adults, will quadruple, affecting 1 in 85 people worldwide (SI 14), fostering tremendous research interest to understand the development and progression of AD. The Alzheimer's Disease Neuroimaging Initiative (ADNI) was launched in 2003 in order to address these research challenges. ADNI's goal is to combine clinical imaging technologies and neuropsychological assessments, along with genetic and other biological data, to study and measure the progression of cognitive impairment into early diagnosis of AD. In 2012, high-coverage (>40x average coverage) whole genome sequencing (WGS) on DNA derived from whole blood of 818 ADNI subjects was performed, in order to assess how genomic variants might be contributing to progression toward, and into, AD.

Many studies have identified a variety of single nucleotide variants (SNVs) throughout the genome that are significantly associated with Alzheimer's Disease (SI 15). There has also been interest in analysis of copy number variants, in particular deletions, in regards to their association to AD susceptibility. Genomic deletions (the loss of genetic sequences on loci scattered across the genome) provide a layer of interpretation often not available with SNVs: the loss of function (LoF) of the underlying sequence that a deletion removes (in a heterozygous or homozygous manner). This additional layer of interpretation is especially important in noncoding regulatory regions of the genome, where there is no obvious way to identify LoF SNVs as there is in coding regions. However, most of the previous studies have only been able to examine very long deletion events (>100 kilo-base-pairs) (SI 16), because of the technologies available, thereby missing deletion events of potential association to AD, especially in noncoding genomic regions (for which single nucleotide polymorphism array probes are less dense compared to genic regions) where the majority of AD genome wide association study (GWAS) signals are located (SI 17). Now, with high-coverage WGS data on ADNI individuals, and the development of higher-resolution deletion algorithms, dramatically increased sensitivity to identify shorter deletion variants is possible.

#### Note S3c

##### ***Data Processing***

*Whole genome sequencing and alignment*

Whole genome sequencing (WGS) on DNA derived from whole blood of 818 ADNI subjects was performed by Illumina's laboratory in 2012-2013, using Illumina HiSeq sequencers, generating 100 base-pair paired-end sequence reads. In 2014, the Broad Institute donated resources to take the Illumina-generated BAM files and re-process the data using Broad's 'best practices' GATK pipeline (SI 18, SI 19, SI 20), with the goal to create improved SNP and indel call accuracy over that generated from Illumina's CASAVA software. Starting from recovered FASTQ files, the sequence reads were mapped to the human reference genome (GRCh37) using BWA-MEM (SI 21), and then processed using the GATK pipeline version GenomeAnalysisTK-3.1-144-g00f68a3.jar), resulting in analysis-ready BAM files. Of the 818 ADNI subjects, 9 were deemed to have provided insufficient consent, and 1 was dropped due to quality control issues during re-processing, leaving 808 subjects on which analysis of their WGS data was subsequently performed. Average genome-wide coverage across the 808 BAM files was ~42x (median: ~41x, range: 33x-81x). The storage size of the 808 BAM files on the disk drive was approximately 150 terabytes.

*ADNI Cohort*

Data used in the preparation of this article were obtained from the Alzheimer's Disease Neuroimaging Initiative (ADNI) database (adni.loni.usc.edu). The ADNI was launched in 2003 as a public-private partnership, led by Principal Investigator Michael W. Weiner, MD. The primary goal of ADNI has been to test whether serial magnetic resonance imaging (MRI), positron emission tomography (PET), other biological markers, and clinical and neuropsychological assessment can be combined to measure the progression of mild cognitive impairment (MCI) and early Alzheimer's disease (AD). For up-to-date information, see www.adni-info.org. The ADNI has been a highly successful and collaborative community effort (SI 22, SI 23), with ADNI data being utilized in approximately 1,800 scientific publications as of January 2020.

*SNV and indel variant discovery*

Broad Institute's 'best practices' GATK pipeline (SI 18, SI 19, SI 20) (version GenomeAnalysisTK-3.1-144-g00f68a3.jar) was run to generate raw single nucleotide variant (SNV) and insertion-deletion (indel) calls. GATK HaplotypeCaller was run on each BAM sample separately, producing single-sample gVCF files. These files were merged into a single gVCF file (using GATK CombineGVCFs). Joint genotyping was then performed (using GATK GenotypeGVCFs) across all 808 samples to produce variant calls. These calls were subsequently filtered (using GATK VariantRecalibrator and GATK ApplyRecalibration) to take into account both sensitivity and specificity. Variant calls that failed the 'Variant Quality Score Recalibration' step of the GATK pipeline were excluded, and genotypes with a genotype quality (GQ) score of <= 20 were set to missing. None of the individuals had a SNV genotype missing rate greater than 1.54%. These procedures produced a total set of 44,535,780 non-monomorphic sequence variants (38,443,567 SNVs and 6,092,213 indels).

*Genotype concordance with microarrays*

Illumina Omni 2.5M microarray data were previously generated on the same ADNI individuals in 2013. We compared the 'PASS' SNVs found in WGS sequencing with those previously called using microarrays using the PLINK software suite (SI 24). We examined biallelic non-monomorphic SNVs using only SNVs whose strands could be matched (including by using the '--flip' option), while excluding SNVs that had the same position but different reference SNV cluster (rs) IDs and excluding SNVs with a missing rate >10%. Only non-missing genotypes in both datasets were considered. Genotype concordance analysis was performed on a total of 1,987,307 SNVs. All of the 808 ADNI samples had at least 99.88% genotype concordance (average of 99.95%) between SNVs found in WGS and those found in the microarray data.

*Identity by descent (IBD) analysis*

To identify any close genetically-related individuals amongst the 808 samples, identity by descent (IBD) estimates called $\hat{\pi}$ were calculated on bi-allelic SNVs using PLINK. To ensure only high-quality genotypes were used in estimating $\hat{\pi}$, stringent quality control filters using PLINK options were employed. These included removal of SNVs with: a) Hardy-Weinberg Equilibrium (HWE) p-value (using the '--hwe' option) of <1 x 10-5 for common variants (defined as SNVs with minor allele frequency (MAF) ≥ 5%) and <1 x 10-2 for rare variants (defined as SNVs with MAF < 5%); b) genotype missing rate of >0.5% (using the '--geno' option); c) genotype concordance rate to microarray data of <99%. Additionally, we performed linkage disequilibrium (LD)-based SNV pruning to obtain a set of independent common SNVs. This procedure involved a) removal of SNVs with MAF <15%; b) application of variance inflation factor (VIF)-based LD-pruning using the '--indep 200 5 1.15' option; and c) application of pairwise genotypic correlation-based LD-pruning using the '--indep-pairwise 100 5 0.1' option. Altogether, after the quality control filters and LD-pruning procedures, 54,210 SNVs were used to compute $\hat{\pi}$. Five pairs of samples were identified that had $\hat{\pi}$ between 0.4-0.6, indicating first-degree relatives. There were no pairs of samples with $\hat{\pi}$ between 0.2-0.4 (second-degree relatives).

*Principal components analysis (PCA)*

To perform a principal components analysis on the WGS ADNI data, the EIGENSTRAT algorithm (SI 25) was used along with data from the 1000 Genomes Project (1000GP) as a reference panel (1000G Phase I v3 ShapeIt2 Reference; 2013-09 haplotype; http://csg.sph.umich.edu/abecasis/MACH/download/1000G.2013-09.html) (SI 26). We performed the previously discussed IBD analysis on the 1000GP reference panel dataset and identified 65 individuals with $\hat{\pi}$>0.2, indicating relatedness of at-least second-degree. These 1000GP individuals and one random individual of each pair of related ADNI samples were dropped from the PCA analysis. To obtain high-quality SNVs for the PCA, the same stringent quality control filters were used as in the IBD analysis, except that variants with <5% MAF were removed (rather than <15% MAF). The 1000GP and ADNI datasets were then merged using overlapping variants, while removing SNVs with inconsistent strands. Additionally, the same LD-based SNV pruning procedures were performed as in the IBD analysis. Altogether, a set of 92,000 independent SNVs were given as input for EIGENSTRAT to compute principal components. Individuals having a PC1 of <-0.01 were considered to be European ancestry (EU) individuals. Out of the 803 ADNI individuals analyzed, 752 were deemed to be EU (~94%). Among the 752 subjects, roughly 31% (233) were classified as brain-normal controls, roughly 45% (342) were classified as exhibiting mild cognitive impairment (MCI), and roughly 24% (177) were classified as exhibiting Alzheimer's Disease (AD) (see Table S3g1 [SI Note S3g]).

*Deletion variant discovery*

To identify deletion variants (>500 base-pairs in length) in the ADNI dataset, the published software algorithm GenomeSTRiP (SI 10) version 1.04.1456 was run on the 808 ADNI re-processed WGS BAM files. GenomeSTRiP combines three lines of technical sequence evidence for calling deletion candidates: breakpoint-spanning reads (split reads), abnormal read-pair separation, and local variation in read depth of coverage, and was previously found to be superior compared to other callers in terms of call specificity, sensitivity, and genotype accuracy (SI 10). Deletions were called with respect to the GRCh37/hg19 version of the human reference genome. The reference genome file used was human_g1k_v37.fasta (downloaded from the 1000 Genomes FTP server (ftp://ftp.1000genomes.ebi.ac.uk/vol1/ftp//technical/reference/human_g1k_v37.fasta.gz). Several genome annotation files are necessary in order for GenomeSTRiP to properly infer genomic deletions from the BAM file data; files compatible with the reference version used were downloaded from the 'svtoolkit' FTP server hosted by the Broad Institute. These included: the genome mask (ftp://ftp.broadinstitute.org/pub/svtoolkit//svmasks/human_g1k_v37.mask.100.fasta.gz), the genome ploidy map (ftp://ftp.broadinstitute.org/pub/svtoolkit//ploidymaps/humgen_g1k_v37_ploidy.map), and the genome copy number mask (ftp://ftp.broadinstitute.org/pub/svtoolkit//cn2masks/cn2_mask_g1k_v37.fasta.gz).

GenomeSTRiP deletion detection consisted of three main workflow phases: pre-processing, discovery, and genotyping. Alternative allele alignment was not performed because the underlying data were 100 base-pair high-coverage WGS BAM files. Default GenomeSTRiP parameters were used. In the pre-processing phase, each ADNI sample was individually processed. BAM file metadata compiled by GenomeSTRiP for each sample was merged for all individuals before the discovery phase. The insert size distribution metadata was merged for all individuals using the 'org.broadinstitute.sv.apps.MergeInsertSizeDistributions' module of GenomeSTRiP. In the discovery phase, all ADNI samples were jointly processed. Minimum and maximum deletion settings were set to 100 and 1,000,000, respectively (using the -minimumSize and -maximumSize options, respectively). To speed GenomeSTRiP computational run time due to the size of the ADNI samples on disk, samples were parallel processed in 5-10 megabase-pair windows (using the -L option), using 1 megabase-pair overlapping windows. The overlapping discovery phase VCF files were merged removing duplicate records using the VCFtools version 0.1.15 'vcf-merge' option (SI 27) and Tabix from htslib version 1.3.2 (http://www.htslib.org/doc/tabix.html). In the genotyping phase, 'PASS' variants from the discovery phase were genotyped across all ADNI samples jointly.

*Deletion QC and filtering*

Deletions were discovered and genotyped by GenomeSTRiP in all 808 ADNI samples, however, to ensure robust computational quality control (access to original DNA samples not being feasible) and downstream population genetic analysis and inference that relies on limited population demographic parameters, only deletions genotyped in individuals deemed to be of 'European ancestry' (752/808, ~94%) (see SI Note S3c, section: Principal components analysis [PCA]) were selected for further analysis. Additionally, because of population genetic forces potentially differing on the X chromosome necessitating special quality control that would be less reliable with only computational tools, and no Y chromosome calls being generated, only autosomal deletions were selected for further analysis. We additionally removed all deletions that were monomorphic (allele frequency=1), indicating reference genome artifacts.

The remaining deletion calls were then individually screened for properties to compile a high-confidence callset (see Fig. S3g1 [SI Note S3g]). Criteria were manually set to maximize the total number of deletion calls while also ensuring reasonable quality control of the accuracy in terms of genotypes and genomic coordinates. To ensure high-quality genotyping of the population at each deletion site, deletions were only retained that had a phred-based genotype quality (GQ) score for all 752 individuals of >=13 (corresponding to ~95% estimated genotype accuracy). Most individuals in most deletions had a reported GQ of 99.

Deletions in individuals either heterozygous or homozygous for the deletion would be expected to have loss of heterozygosity (LOH) in the portion of their chromosome where the deletion resides. Therefore, SNV concordance was measured in relevant individuals within deletion coordinates. Deletions were only retained where all individuals had <=25% SNV discordance. Discordance here is defined as the proportion of SNVs that are discordant in any relevant het-del or hom-del individual versus expectation from a true deletion call using GATK-derived SNV calls as the assumed 'gold-standard' correct genotype. Having a high SNV discordance in an individual would indicate that either that individual was not well genotyped for the deletion or that the coordinates of the deletion are largely misspecified. Deletions for which there were no overlapping SNVs were additionally retained.

Because high-quality deletion coordinate localization is important for biological interpretation, only deletions with breakpoint (start coordinate and end coordinate) confidence intervals (as given from GenomeSTRiP output) corresponding to <=3% of deletion length were retained. We use 'average' deletion coordinates to represent final coordinates (not extremes) since we don't want to induce false genomic annotation overlaps in downstream analysis.

Because of the way GenomeSTRiP genotypes candidate deletions, it is possible for a larger deletion overlapping a smaller deletion to be simultaneously genotyped in the same individual. This would be the case, for example, when a longer common-frequency deletion overlaps a shorter deletion that is a singleton/rare-frequency deletion in another individual(s). In situations like this, rule-based filters were used to clarify genotype assignment and collapse redundant calls. Filter rules were applied as follows:

-Deletions which extend further in both start and end directions will have corresponding redundant genotypes in shorter, fully-overlapped deletion candidates. Shorter deletion candidates are collapsed into larger deletions for all matching genotypes. Remaining shorter deletions with now mutually exclusive genotypes are kept as correct calls.
-Overlapping deletions with mutually exclusive genotypes are deemed to be separate deletions.
-If two deletions are completely overlapped or have very close coordinates (>90% overlapping), the deletion genotype with homozygous deletion calls is deemed to be the correct genotype and the other coordinates are ignored.
-Two deletions with very close coordinates (>90% overlapping) that share all genotypes, plus a few additional for one of the deletions, is deemed to be a single deletion event. Since the deletion is likely real and common, the deletion call with the most genotyped individuals is deemed to be the correct deletion genotype and coordinates.
-When two deletions have less than 80% overlapping coordinates, heterozygous and homozygous deletion genotypes are seen as coming from different deletion events (such as when one deletion is common in allele frequency and the other deletion is a singleton).
-When a rare, longer deletion overlaps a rare, shorter deletion, any genotypes shared between the deletions is assigned to the longer deletion since GenomeSTRiP would likely have also assigned the genotype(s) to the shorter one because of the overwhelming technical support given from the longer one.
-When two deletions of similar length have very close overlapping coordinates (>90%), if a singleton/rare call has genotypes also present in the other common deletion, the genotypes for the singleton/rare deletion are retained and removed from the other deletion. In this type of situation, it is likely that GenomeSTRiP added the same genotypes to the common deletion because of the nature of the joint-calling algorithm which may in some circumstances give more weight to common alleles.
-Some deletions cannot be interpreted in light of ambiguous genotyping, such as when one individual genotype is shared between two partially overlapping deletions. When genotypes are not able to be confidently assigned using these filter rules, the corresponding deletions are dropped from further analysis.

Deletions remaining after all prior QC and filtering steps were assessed for violation of Hardy-Weinberg Equilibrium (HWE) (using the '--hwe' option of VCFtools [SI 27]) in order to identify deletions undergoing obvious selection pressures other than purifying natural selection, or to identify deletions with low-quality genotyping. The threshold of removal was set to a HWE p-value <=1x10-5. Of the remaining deletions, only 0.3% were between the range of 1x10-2 < p < 1x10-5. After application of all QC criteria and filtering steps, a set of 10,619 autosomal deletions genotyped within the 752 ADNI EU individuals remained.

#### Note S3d

***Data Records***

We deposited [in-progress] four data records to the Laboratory of Neuro Imaging (LONI) Image and Data Archive (IDA) hosted at the University of Southern California, Los Angeles, CA, USA: the original GenomeSTRiP deletion callset VCF file-before quality control and filters were applied, the quality controlled and filtered deletion callset with individual genotypes-VCF file, the quality controlled and filtered deletion callset with summary counts for each phenotype-BED file, and the quality controlled and filtered deletion callset with summary population allele frequency to the LONI IDA. The original WGS BAM files are available via hard drives from the LONI IDA. The re-aligned WGS sequence data using GATK-best practices were processed at the Broad Institute on live disk storage, but due to the size of the data, were subsequently moved to 'cold' storage offsite. These re-aligned data may no longer be available given the cost of data maintenance of ~150 terabytes. The SNV/indel callset was previously posted to the LONI IDA. All files obtained from the LONI IDA require that investigators download, review, sign, and submit the ADNI WGS Data Use Agreement and be a registered user of ADNI data. More information on obtaining data access can be found at: http://adni.loni.usc.edu/data-samples/access-data/ .

#### Note S3e

##### ***Technical Validation***

Previous validation has been extensive for the GATK pipeline, widely used in the genomics sequencing community, as well as for the deletion-calling algorithm GenomeSTRiP, employed in multiple consortium efforts including the 1000 Genomes Project (SI 6). GenomeSTRiP has been previously found to offer advantages in both sensitivity and specificity in comparison with many other deletion callers (SI 10). Therefore, we focused our validation efforts on examining the properties of the distributions of sequence variants that we generated, expecting high-quality distributions, qualitatively similar to that observed in previous successful studies using these tools.

*Re-processed BAM file integrity*

To ensure the BAM file re-processing procedure was performed at high-quality for the 808 samples, SNV variants discovered in our callset were compared with SNVs identified on microarray data previously performed on the same ADNI subjects (see SI Note S3c, section Genotype concordance with microarrays). Genotype concordance between the WGS SNV callset and the SNV microarray data was very strong with an average genotype concordance of 99.95% (minimum 99.88%). Additionally, the transition/transversion (Ti/Tv) ratio of 2.02 for novel SNVs discovered in our callset was comparable with the Ti/Tv ratio of ~2.2 for SNVs catalogued in dbSNP (https://www.ncbi.nlm.nih.gov/snp/). Also, genic SNVs annotated using MapSNPs (part of the PolyPhen-2 software suite (SI 5) resulted in a Ti/Tv ratio for coding SNVs of 2.97, comparable with findings in other datasets of ~3 (SI 28).

*Allele frequency spectrum shape*

Deletion mutations initially start as a singleton in allele frequency in the population, deriving from a de-novo mutational event in the germline. Deletion mutation recurrence at the same start and end coordinates is extremely unlikely due to chance because of low mutation rates (SI 29). Therefore, in considering the shape of the distribution of deletion variant allele frequencies in a population, most events are rare, except a few events that occurred many generations ago (and are therefore present in high frequency across worldwide or broad demographic populations), or have arisen to high frequency due to positive natural selection in favor of the deletion allele. However, since many deletions overlap at minimum a functional regulatory element (especially for deletions >10,000 base-pairs), widespread positive selection on deletions is not observed; conversely, negative selection is observed (SI 6). To assess whether our deletion callset matches qualitative expectations in the shape of the allele frequency spectrum, we rank order all deletions by genotype frequency. Fig. S3g2 (SI Note S3g) shows a cumulative fraction plot of the deletion allele frequency across the deletion callset. The smooth curve of the ranking shows the expected pattern of abundant distinct rare variant counts transitioning into infrequent common variant counts (SI 6). The abundance of distinct rare variants in our callset (~75% of our deletion calls are tripleton or lower in allele frequency) is likely at least partially due to enhanced sensitivity in deletion calling available with the high-coverage ADNI WGS data.

*Common deletion variant concordance*

Rare variant deletion calls, often unique to one particular dataset, are difficult to validate as true calls (some may instead be false positives due to technical artifacts in the underlying data) without experimental evidence in the genotyped samples. However, common deletion variants should persist across datasets when the underlying population samples are from the same broad demographic population. The 1000 Genomes Project (1000GP) consortium has released a set of breakpoint-resolved deletion calls (SI 6, supplemental table 3]), derived from a combination of multiple structural variant callers used in variant discovery and genotyping including GenomeSTRiP, as a part of their goal to characterize common genomic variation in worldwide populations. Taking deletions found to be >=5% allele frequency in the EUR population (broadly of European ancestry) from the 1000GP callset (SI Note S2a), and comparing with ADNI deletions >=1% allele frequency from our quality-controlled and filtered callset, we find that for ADNI deletions greater than or equal to our median call length (3,114 base-pairs), there is a 92.1% (279 ADNI/303 1000GP) concordance rate at a minimum of 90% overlapping coordinates. This means that for deletions at our median length or longer, we have both high sensitivity and high breakpoint accuracy to correctly identify deletion variants. However, we do note that for shorter lengths below our median length, we observe deletion sensitivity loss: e.g. there is a 55.6% confirmation rate for ADNI deletions >=1,000 base-pairs at a minimum of 90% overlapping coordinates (this analysis assumes that the 1000GP calls are the correct calls). As with all structural variation callsets (which can have sensitivity loss due to the short-read sequencing technology typically employed, as was the case with the ADNI WGS), the absence of a deletion variant at a particular locus does not indicate that the deletion is not present in the population, rather it indicates that at the particular QC and filter thresholds used (such as by GenomeSTRiP and our downstream workflow), a deletion variant could not be called with statistical confidence. It is likely that at least a portion of this sensitivity loss (compared with 1000GP calls) at shorter deletion lengths arises from deletion calls failing our QC and filtering protocol, as well as deletion calls originating from a different deletion algorithm other than GenomeSTRiP used by the 1000GP in the compilation of their dataset.

*Flat quantile-quantile plot*

Since the ADNI WGS dataset is likely underpowered from the perspective of identifying common variants with large effect size on the phenotype of interest, consistent with prior association analyses from AD-GWAS (SI 16, SI 17), (our dataset may be useful instead for targeted analyses, see SI Note S3f), we would expect that a case-control analysis of common variant loci would result in only a few loci that would exhibit at most marginal significance. This case-control comparison can be represented using a quantile-quantile (Q-Q) plot. If the realized distribution of p-values from the actual case-control comparisons is similar to the null expectation distribution of p-values, then the points in the Q-Q plot will lie approximately on the diagonal line, corresponding to y = x. Using Fisher's exact-test p-values of deletion counts between cases and controls at each common variant locus, Fig. S3g3 (SI Note S3g) shows the resulting Q-Q plots, for the three possible phenotype groupings (Control vs AD, Control+MCI vs AD, Control vs MCI+AD). The most significant single locus occurs in the Control vs MCI+AD phenotype grouping comparison, with original p-value of 0.00031427. However, when applying multiple-test correction to the result, the corrected p-value is 0.43, in line with expectations from an underpowered dataset. The flatness of the Q-Q plots ('well-behaved'), compared to null expectation, indicates high-quality genotyping across subsets of the ADNI deletion callset. This enables confidence in the accuracy of the genotyping in the callset across the population as a whole, which is useful in other research contexts (see SI Note S3f, section: Usage case 3: Population deletion callset), since the vast majority of deletions in the callset would not be phenotype-specific.

#### Note S3f

***Usage Cases***

*Usage case 1: LoF analysis in combination with SNVs and indels*

Detecting loss of function (LoF) in the genome from genetic mutations can be difficult, especially in noncoding regions; however, indels (defined as <=50bp) and deletions allow inference of loss of regular biological function (which may sometimes technically be gain of immediate function if the deleted region encodes a suppressor, etc). Joint analysis of indels and deletions may uncover regions of the genome where an individual or group of individuals is homozygous for loss of function at a particular locus (heterozygous for a deletion indel on one chromosome and heterozygous for a deletion on the other chromosome). Additional analysis involving SNVs in combination with indels and/or deletions may also uncover interesting biological activity at loci where these 'complex' heterozygotes occur, due to joint interactions from variants on each parental chromosome.

*Usage case 2: Deletion variant network burden testing*

While many cohort-based datasets fail to uncover single variants of large effect, there is tremendous interest in the medical genetics community to understand how the collective burden of a group of variants may be contributing together across a population to influence susceptibility toward the phenotype of interest, or affect the severity (penetrance) of the trait (SI 30). Deletions present in this dataset are on average very rare (75% of all deletions in the callset have an allele frequency of tripleton or lower). These rare variant calls may provide insight into the genetic etiology of Alzheimer's Disease, when taken together as hypothesis-driven sets of related biological units. For example, a set of deletions overlapping genes highly expressed in relevant tissue/cell types, or a set of noncoding deletions overlapping important regulators (such as enhancers or DNase I hypersensitive sites) in relevant tissues/cell types, or other similarly formed sets based upon biological intuition, may provide insight into the contribution of rare variants towards onset of Alzheimer's Disease. This network burden analysis can be especially useful in noncoding regions where the presence of a deletion can be seen as a heterozygous loss of function (see also SI Note S3f, section: Usage case 1: LoF analysis in combination with SNVs and indels), and so a set of deletions could be assembled to assess the collective burden imposed by variants overlapping regulatory loci of interest.

*Usage case 3: Population deletion callset*

The vast majority of the deletions identified in this callset would not be related to any particular phenotype, but instead represent a collection of segregating genomic variants identified in an otherwise healthy population. This is especially the case given that Alzheimer's Disease is a post-reproductive phenotype (all samples were collected from individuals that were at least 50 years old) and selection might therefore be modest or absent for any AD-related variants in the dataset, unless pleiotropic in nature. There is great interest in studying large collections of individuals for analysis of segregating genomic variation to learn about mutational processes and natural selection, amongst other fundamental biological and evolutionary processes. Since this deletion callset was generated from only individuals of European ancestry (see SI Note S3c, section: Principal components analysis [PCA]) and also has properties of a robust population dataset (see SI Note S3e), the deletion calls included in this dataset (especially the deletions at the median length or longer, where sensitivity was greatest) could be of use in projects that require these properties for deletion variant analysis.

#### Note S3g

**Table S3g1. European subject cohort within ADNI.**

Of the 808 ADNI subjects for which we analyzed whole genome sequencing data, 752 subjects were determined to be of European ancestry using principal components analysis. 'Control' phenotype corresponds to brain-healthy cognition. 'MCI' phenotype corresponds to mild-cognitive impairment. 'AD' phenotype corresponds to Alzheimer's Disease diagnosis. Deletions genotyped within these individuals were selected for further downstream analysis.

| Phenotype | Control | MCI | AD |
|---|---|---|---|
| Number of Subjects | 233 | 342 | 177 |
| Percentage of Cohort | 31 | 45 | 24 |

**Table S3g2. ADNI Deletion callset characteristics.**

Deletion allele frequency and length characteristics of the quality-controlled and filtered final deletion callset among 752 European-ancestry ADNI individuals. The abbreviation 'bp' corresponds to DNA base-pairs.

| Deletion Callset Characteristic | |
|---|---|
| Number of Deletions | 10,619 (100%) |
| Singleton AF | 6,443 (60.7%) |
| Doubleton AF | 1,051 (9.9%) |
| Tripleton AF | 483 (4.5%) |
| >=1% AF | 1,366 (12.9%) |
| Average Length | 9,285 bp |
| Median Length | 3,114 bp |
| Minimum Length | 440 bp |
| Maximum Length | 853,585 bp |
| Average Length Singleton AF | 10,735 bp |
| Median Length Singleton AF | 3,244 bp |
| Average Length >=1% AF | 6,155 bp |
| Median Length >=1% AF | 3,367 bp |

**Figure S3g1: Schematic overview of the ADNI deletion callset generation.**

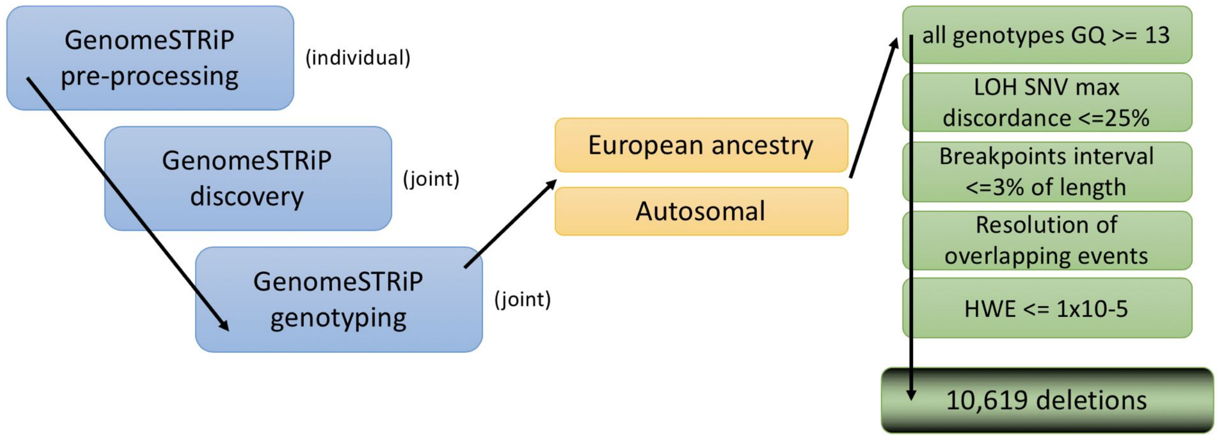


**Figure S3g2: Cumulative fraction of ADNI deletion allele frequency.**

Deletions in the final quality-controlled and filtered callset are rank-ordered from lowest allele frequency to highest. The color gradient slowly changing from dark green to light grey corresponds to different observed allele frequency value levels throughout the dataset.

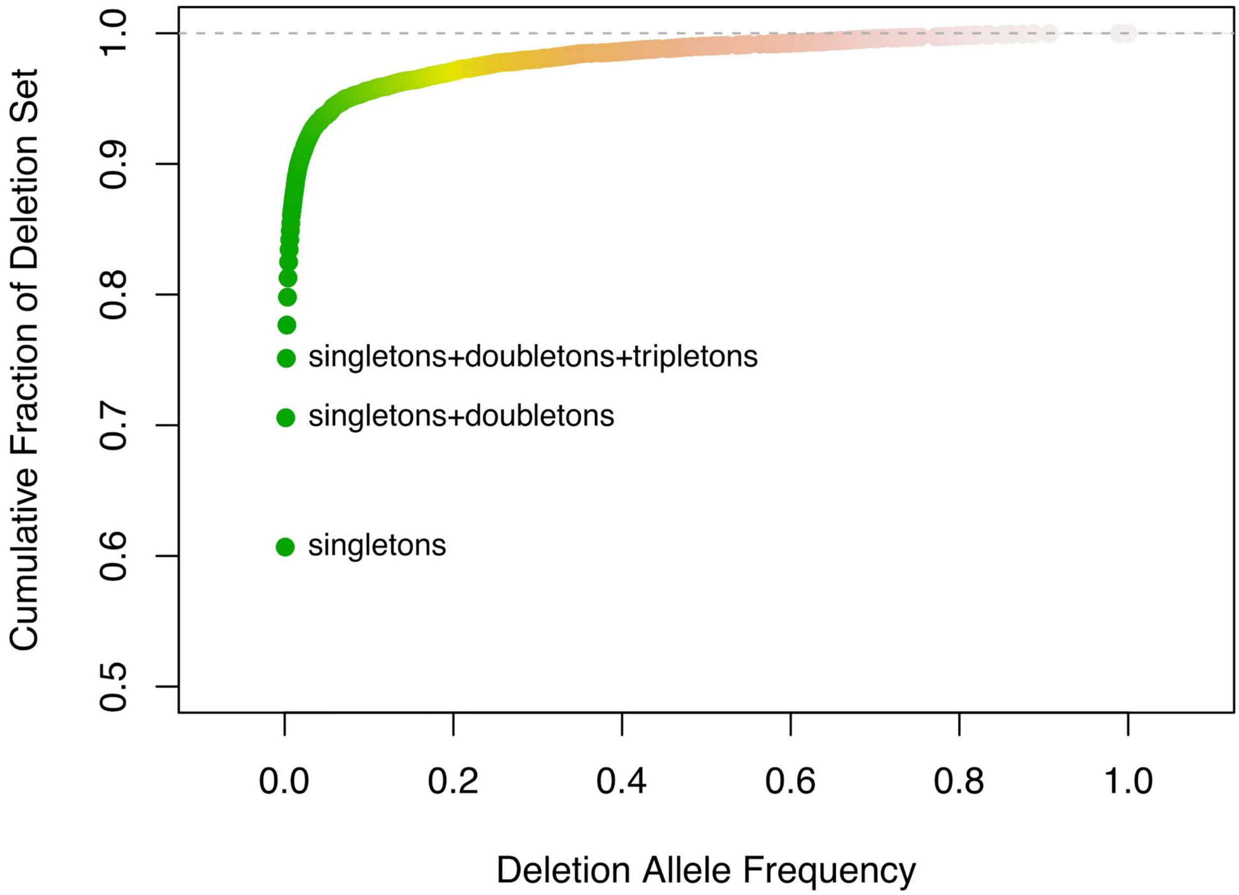

**Figure S3g3: Q-Q plots of phenotype comparisons.**

Quantile-quantile plots of the quality-controlled and filtered deletion callset, before Hardy-Weinberg Equilibrium filter. P-values of each deletion's association with the phenotype ('Control'=brain-healthy cognition, 'MCI'=mild cognitive impairment, 'AD'=Alzheimer's Disease) were determined from Fisher's exact test. Since it is unclear if MCI better associates with Control or with AD, in **a)** MCI is not included, in **b)** Control+MCI are merged, and in **c)** MCI+AD are merged.

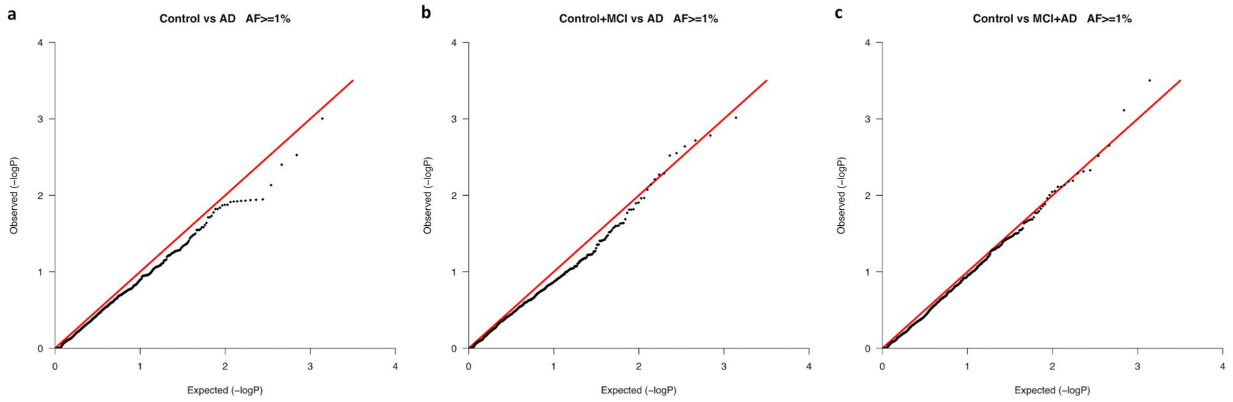


### Note S4

**Regulatory datasets**

#### Note S4a

***NIH Roadmap Epigenomics Consortium (REC)***

Regulatory element chromatin accessibility data (DNase I hypersensitivity) and histone modification data (H3K4me1 "enhancer", H3K36me3 "transcribed", H3K27me3 "polycomb-repressed", H3K9me3 "heterochromatin") were downloaded from the supplementary website for the 2015 NIH Roadmap Epigenomics Consortium ("REC") paper (SI 1) located at https://egg2.wustl.edu/roadmap/web_portal/index.html. All data we used were derived from REC 'consolidated epigenomes', which were uniformly reprocessed and standardized epigenomes, designed to eliminate differences between research centers and changes to sequencing technology that occurred over the course of the REC project. Two types of DNase I callsets with base-pair resolution were analyzed: calls made from the Hotspot algorithm (SI 3) which uses a binomial distribution model (from https://egg2.wustl.edu/roadmap/data/byFileType/peaks/consolidated/broadPeak/), and calls made from the MACS algorithm (SI 2) which uses a Poisson distribution model (from https://egg2.wustl.edu/roadmap/data/byFileType/peaks/consolidated/narrowPeak/). Both callsets that were generated with an expected false discovery rate (FDR) of 1% were chosen. We used DNase I callsets from the two different algorithms to assess consistency between the results. For histone modification data, callsets with base-pair resolution made from MACS with a 1% expected FDR were chosen (from https://egg2.wustl.edu/roadmap/data/byFileType/peaks/consolidated/narrowPeak/).

To ensure reliable regulatory data in the analysis of overlapping genomic deletions, we selected only tissues/cell-types for analysis that were primary in nature, i.e. not embryonic stem (ES) cells, induced pluripotent stem (iPS) cells, or ES-derived cells, as non-primary tissues/cell-types have undergone significant passaging effects from which structural variant artifacts could accumulate in the cells (SI 31, SI 32). We selected all primary tissues/cell-types which were available across both types of DNase I callsets and all four histone modification callsets. We excluded two cell lines, E017 and E124, that appeared to be highly similar (same tissue/cell-type and same relative donor age) to E088 and E029, which were retained. Of the 25 tissues/cell-types selected, 15 were derived from fetal samples, and 10 were derived from adult (three years or older) samples. SI Table S3 summarizes the selected tissues/cell-types. These data assume that regulatory elements are shared across all human populations, as the assays performed included samples from various ancestries.

#### Note S4b

##### ***TAD-loops***

Topologically associated domain loop (TAD-loop) boundary data were downloaded from GEO accession GSE63525 (from https://www.ncbi.nlm.nih.gov/geo/query/acc.cgi?acc=GSE63525), data generated using in situ Hi-C from Rao & Huntley et al. (SI 33). Files with names ending in "*_HiCCUPS_looplist_with_motifs.txt.gz" were selected and the genomic coordinates corresponding to TAD-loop boundaries were extracted. As with the DNase I hypersensitivity and histone modification callsets data, only primary tissues/cell-types were selected for analysis. The five callsets selected were derived from GM12878, NHEK, IMR90, HUVEC, and HMEC tissues/cell-types.

#### Note S4c

##### ***ENCODE Uniform CTCF TF Peaks***

To overlay CTCF transcription factor binding sites within TAD-loops, we used CTCF locations from the same tissues/cell-types examined for TAD-loops. CTCF ChiP-seq callset data, processed in a uniform pipeline from the ENCODE project March 2012 data freeze (SI 34), were downloaded from the UCSC genome browser "Transcription Factor ChIP-seq Uniform Peaks from ENCODE/Analysis" track at http://genome.ucsc.edu/cgi-bin/hgFileUi?db=hg19&g=wgEncodeAwgTfbsUniform. Each tissue/cell-type had more than one callset, each originating from a different analysis center. For each tissue/cell-type separately, all CTCF calls across files from the various analysis centers were merged using the BEDTools (SI 12) 'merge' option. Then, only CTCF sites from a given tissue/cell-type which overlapped TAD-loops in the same tissue/cell-type were extracted and retained. In this way, mis-specification in either TAD-loop or CTCF callsets are then not contaminated between analyses. This is especially important in pleiotropic analyses where concordance between a particular CTCF and a particular TAD-loop are assumed to co-occur in the same tissue(s)/cell-type(s). This limits counting transient CTCF binding sites across the tissues/cell-types.

### Note S5

**Deletion Simulations**

#### Note S5a

***Deletion simulation strategy***

Deletion length can be a confounder in analyses of deletions overlapping regulatory features and associated cellular pleiotropy measures, because longer deletions have a greater chance than shorter deletions to randomly hit sparse annotations in the genome (such as regulatory elements, especially cellularly pleiotropic regulatory elements). Most overlap statistics that could be computed (such as binary association, bp_affected [SI Note S1a], or PlyRS measures [SI Note S1c]) would then have an inherent length bias, making analyses confounded. Additionally, longer deletions are typically easier for deletion callers to identify because missing sequence coverage over a longer length appears more statistically significant. Therefore, to ensure reliable interpretation of deletion overlap within regulatory regions in a length-controlled manner, we developed a simulation strategy to place mock deletion copies (of the same length, and retaining the AF label for downstream analysis) randomly along the genome (using only allowable genomic space [SI Not S2c]), but keeping the mock copy on the same chromosome (but not the same chromosome arm/locus band as might introduce non-independence of the simulations) and same genomic compartment space (intronic or intergenic), to approximate local context-dependent effects. For each deletion, we created 1,000 mock deletion copies, thereby creating 1,000 mock deletion datasets each with a random distribution of deletion locations. To later compare significance of overlap associations, we created an additional 1,000 mock deletion copies for each deletion (additional 1,000 mock deletion datasets). Using this length-matched simulation framework, we are able to analyze both horizontal and vertical axes on which purifying selection may be operating on deletions (SI Note S1). We randomized deletions, as opposed to regulatory elements, utilizing the fact that the deletions are mutations (relative to the human reference genome) while assuming that regulatory elements are essentially fixed (i.e. consistent) components in the modern human genome across populations.

#### Note S5b

***Simulation significance calculation***

We have developed the following procedure to detect reduction in deletion variation. For each real (i.e. observed) deletion, we compare, one at a time, a measure of interest (e.g. $PlyRS_{max}$ [SI Note S1c]) of the real deletion's value, to each mock deletion copy's (i.e. expected) value of the measure of interest. Each time that the real deletion has a lower (or equal) value than a mock copy (which indicates overlap in the real deletion compared to simulation had the same amount or less), we assign that instance to a counter and perform over all 1,000 mock deletion copies. We include the ‘equal to’ so that information from the simulation is utilized, otherwise all deletions with no real overlap will always receive an empirical p-value of 0.001 ([no counts + 1] / 1000), but including the ‘equal to’ means that the number of simulation no-overlaps will be included in the empirical p-value. This information is especially useful when considering deletions with no real overlap of different lengths. At the end of this process, we calculate the one-sided empirical p-value of this analysis, taken as the counter of instances (plus 1 since our real result can be considered an instance of observation unless counter=1000 at which point we ignore our observation) divided by 1,000 tests. Therefore, for each deletion, there is an empirical p-value of that deletion's measure of interest versus 1,000 mock copies. We additionally perform this same process for each of the 1,000 mock deletion sets against 1,000 additional matched mock deletion sets. Therefore, for each original mock deletion set, there is an empirical p-value of every deletion's measure of interest versus that of the 1,000 additional mock copies. To assess significance of the depletion results, we calculate the sum of the natural log(empirical p-value) for every deletion in the real dataset, and additionally calculate this sum for every original mock deletion dataset. Using the distribution of this sum for the mock deletion datasets, we perform a t-test of where the real deletion dataset sum resides amidst the mock distribution. We can use a t-test because the mock distribution from the sum of ln(emp p-val) is approximately normal (see SI Fig. S5b1 for an example). From the t-test, we can also derive the effect size of the result (i.e. Cohen's D, in units of standard deviations), standardizing the interpretation when comparing results across regulatory features (each of which may have been composed of a different sample size of overlapping deletions).

Calculation steps in outline format:

- -Create 1,000 mock deletion copies for each real deletion
- -For each real deletion, compare a measure of interest (e.g. $PlyRS_{max}$) to each deletion copy
- -Each time that the real deletion has a value <= a mock copy value (indicates depletion) assign that instance to a counter
- -Perform over all 1,000 mock copies
- -Calculate the one-sided empirical p-value of this analysis (counter/1,000)
- -Create 1,000 mock deletion copies again
- -Repeat analysis of steps above for each original mock copy compared to its own 1,000 new mock copies
- -Calculate the sum of the ln(emp p-val) of all deletions in the real dataset, and the sum for all deletions in all original 1,000 mock datasets
- -Use the distribution of this sum for the mock datasets to perform a t-test of where the real deletion dataset sum resides
- -Convert this into effect size in units of standard deviation (Cohen's D)

We measure depletion relative to the deletion, not relative to a percentage of a regulatory element, since the exact boundary of an element can be uncertain across multiple tissues/cell-types (*see SI Appendix, Note S1a*). Because longer deletions in our datasets have the ability to potentially overlap an entire distinct regulatory element or even multiple elements, we want to capture that information in our simulation experiments.

**Figure S5b1: Sum of ln(emp p-val) distribution example.**
Sum of logarithm(empirical p-value) from real 1000GP deletion set (red line with its datapoint) compared to simulations for $PlyRS_{max}$. T-test is significant at $5.84x10^{-09}$ indicating that the 1000GP deletion set is depleted of $PlyRS_{max}$ compared to null expectation.

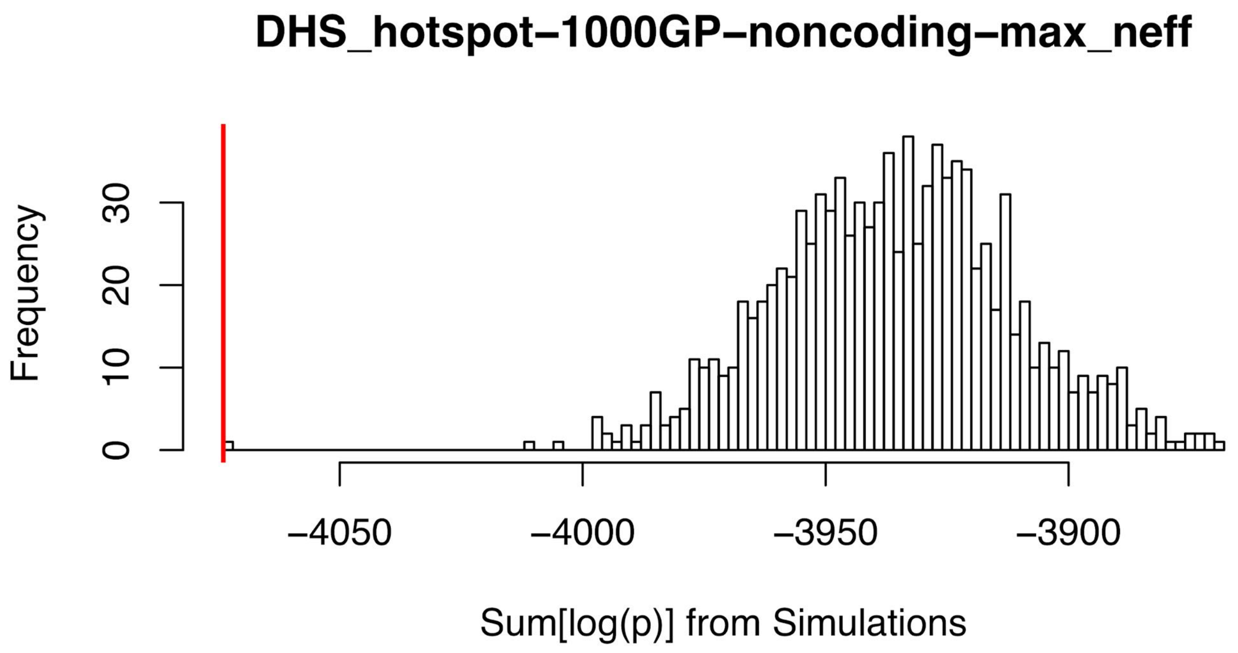


Note S5c

***Simulation space allowed***
For examination of deletions overlapping regions of chromatin accessibility or histone modifications, we exclude deletions and loci within the genome that have TAD-loop annotation (SI Note S4b) while working within the otherwise full genomic coordinates allowed (SI Note S2c-S2d). This is to ensure that any signal of purifying selection observed for chromatin accessibility or histone modifications is not contaminated with co-localized selection due to TAD-loop disruption. Since TAD-loop annotations are sparse in the genome (affecting ~10% of deletions), we analyze TAD-loop depletion using the full genomic coordinates allowed (SI Note S2c-S2d), as excluding TAD-loops with co-localizing chromatin accessibility or histone modifications (which are much shorter in length) would greatly reduce the number of TAD-loops we could analyze, thereby inhibiting robust analysis of potential selection to preserve TAD-loops. Additionally, when subsetting the deletions overlapping TAD-loop into those that overlap TAD-loop annotation but not coincident CTCF annotation, we perform a separate allowable genome space including the full genomic coordinates allowed excluding coordinates with CTCF sites within TAD-loop annotations (SI Note S4c). We do not make a distinction between a TAD-loop annotation for which CTCF sites are excluded within its coordinates and a TAD-loop annotation for which no CTCF site was originally observed. Using this separate allowable genome space for simulations ensures that the depletion values we observe for TAD-loop_noCTCF are reflective of the exclusionary criteria we use when conditioning on the real set of deletions which

also exclude coordinates with CTCF sites. For all analysis involving CTCF, we analyze only CTCF sites that are identified in the same tissue/cell-type as the coincident TAD-loop.

#### Note S5d

**Depletion simulation results**

**Table S5d1. Depletion simulation results for DHS, enhancer, transcribed, polycomb-repressed, or heterochromatin ($PlyRS_{sum}$).**

| regclass | dataset | gencompartment | cohensd | cohensd_lower | cohensd_upper |
|---|---|---|---:|---:|---:|
| DHS_hotspot | 1000GP | noncoding | 6.70 | 6.40 | 7.00 |
| DHS_hotspot | 1000GP | intronic | 5.04 | 4.81 | 5.27 |
| DHS_hotspot | 1000GP | intergenic | 4.31 | 4.11 | 4.51 |
| DHS_hotspot | ADNI | noncoding | 5.45 | 5.21 | 5.70 |
| DHS_hotspot | ADNI | intronic | 3.44 | 3.28 | 3.61 |
| DHS_hotspot | ADNI | intergenic | 4.25 | 4.05 | 4.45 |
| DHS_MACS | 1000GP | noncoding | 7.14 | 6.82 | 7.46 |
| DHS_MACS | 1000GP | intronic | 5.73 | 5.48 | 5.99 |
| DHS_MACS | 1000GP | intergenic | 4.28 | 4.09 | 4.48 |
| DHS_MACS | ADNI | noncoding | 5.49 | 5.25 | 5.74 |
| DHS_MACS | ADNI | intronic | 3.69 | 3.51 | 3.86 |
| DHS_MACS | ADNI | intergenic | 4.08 | 3.89 | 4.27 |
| enhancer_H3K4me1 | 1000GP | noncoding | 4.32 | 4.12 | 4.52 |
| enhancer_H3K4me1 | 1000GP | intronic | 2.54 | 2.41 | 2.67 |
| enhancer_H3K4me1 | 1000GP | intergenic | 3.86 | 3.68 | 4.04 |
| enhancer_H3K4me1 | ADNI | noncoding | 4.87 | 4.65 | 5.09 |
| enhancer_H3K4me1 | ADNI | intronic | 3.06 | 2.91 | 3.21 |
| enhancer_H3K4me1 | ADNI | intergenic | 4.04 | 3.85 | 4.23 |
| transcribed_H3K36me3 | 1000GP | noncoding | -0.70 | -0.77 | -0.63 |
| transcribed_H3K36me3 | 1000GP | intronic | -1.19 | -1.27 | -1.11 |

| | | | | | |
|---|---|---|---|---|---|
| transcribed_H3K36me3 | 1000GP | intergenic | 2.19 | 2.08 | 2.31 |
| transcribed_H3K36me3 | ADNI | noncoding | -1.71 | -1.80 | -1.61 |
| transcribed_H3K36me3 | ADNI | intronic | -1.98 | -2.09 | -1.87 |
| transcribed_H3K36me3 | ADNI | intergenic | 0.69 | 0.62 | 0.76 |
| polycomb_H3K27me3 | 1000GP | noncoding | 0.26 | 0.20 | 0.32 |
| polycomb_H3K27me3 | 1000GP | intronic | 1.58 | 1.49 | 1.67 |
| polycomb_H3K27me3 | 1000GP | intergenic | -0.85 | -0.92 | -0.77 |
| polycomb_H3K27me3 | ADNI | noncoding | 2.18 | 2.06 | 2.29 |
| polycomb_H3K27me3 | ADNI | intronic | 1.36 | 1.27 | 1.44 |
| polycomb_H3K27me3 | ADNI | intergenic | 1.74 | 1.64 | 1.84 |
| heterochromatin_H3K9me3 | 1000GP | noncoding | -1.18 | -1.26 | -1.10 |
| heterochromatin_H3K9me3 | 1000GP | intronic | -1.49 | -1.58 | -1.40 |
| heterochromatin_H3K9me3 | 1000GP | intergenic | -0.48 | -0.55 | -0.42 |
| heterochromatin_H3K9me3 | ADNI | noncoding | -3.40 | -3.56 | -3.24 |
| heterochromatin_H3K9me3 | ADNI | intronic | -2.18 | -2.29 | -2.07 |
| heterochromatin_H3K9me3 | ADNI | intergenic | -2.61 | -2.74 | -2.48 |

**Table S5d2. Depletion simulation results for DHS or enhancer ($PlyRS_{sum\text{-}mono}$).**

| regclass | dataset | gencompartment | cohensd | cohensd_lower | cohensd_upper |
|---|---|---|---|---|---|
| DHS_hotspot | 1000GP | noncoding | 4.46 | 4.25 | 4.66 |
| DHS_hotspot | 1000GP | intronic | 2.95 | 2.81 | 3.10 |
| DHS_hotspot | 1000GP | intergenic | 3.35 | 3.19 | 3.51 |

| | | | | | |
|---|---|---|---|---|---|
| DHS_hotspot | ADNI | noncoding | 4.14 | 3.94 | 4.33 |
| DHS_hotspot | ADNI | intronic | 1.88 | 1.78 | 1.98 |
| DHS_hotspot | ADNI | intergenic | 4.01 | 3.82 | 4.20 |
| DHS_MACS | 1000GP | noncoding | 6.22 | 5.94 | 6.50 |
| DHS_MACS | 1000GP | intronic | 4.75 | 4.53 | 4.96 |
| DHS_MACS | 1000GP | intergenic | 3.91 | 3.73 | 4.09 |
| DHS_MACS | ADNI | noncoding | 4.48 | 4.27 | 4.68 |
| DHS_MACS | ADNI | intronic | 2.96 | 2.82 | 3.10 |
| DHS_MACS | ADNI | intergenic | 3.35 | 3.19 | 3.50 |
| enhancer_H3K4me1 | 1000GP | noncoding | 3.32 | 3.16 | 3.48 |
| enhancer_H3K4me1 | 1000GP | intronic | 2.20 | 2.08 | 2.31 |
| enhancer_H3K4me1 | 1000GP | intergenic | 2.66 | 2.53 | 2.79 |
| enhancer_H3K4me1 | ADNI | noncoding | 4.23 | 4.03 | 4.43 |
| enhancer_H3K4me1 | ADNI | intronic | 2.42 | 2.29 | 2.54 |
| enhancer_H3K4me1 | ADNI | intergenic | 3.66 | 3.49 | 3.84 |
| transcribed_H3K36me3 | 1000GP | noncoding | 0.68 | 0.61 | 0.75 |
| transcribed_H3K36me3 | 1000GP | intronic | 0.19 | 0.13 | 0.25 |
| transcribed_H3K36me3 | 1000GP | intergenic | 1.76 | 1.66 | 1.86 |
| transcribed_H3K36me3 | ADNI | noncoding | -0.56 | -0.63 | -0.50 |
| transcribed_H3K36me3 | ADNI | intronic | -0.71 | -0.78 | -0.64 |
| transcribed_H3K36me3 | ADNI | intergenic | 0.38 | 0.32 | 0.45 |
| polycomb_H3K27me3 | 1000GP | noncoding | 1.46 | 1.37 | 1.54 |
| polycomb_H3K27me3 | 1000GP | intronic | 1.93 | 1.82 | 2.03 |

| | | | | | |
|---|---|---|---|---|---|
| polycomb_H3K27me3 | 1000GP | intergenic | 0.36 | 0.30 | 0.43 |
| polycomb_H3K27me3 | ADNI | noncoding | 2.04 | 1.93 | 2.15 |
| polycomb_H3K27me3 | ADNI | intronic | 1.34 | 1.25 | 1.42 |
| polycomb_H3K27me3 | ADNI | intergenic | 1.58 | 1.48 | 1.67 |
| heterochromatin_H3K9me3 | 1000GP | noncoding | -0.45 | -0.51 | -0.38 |
| heterochromatin_H3K9me3 | 1000GP | intronic | 0.49 | 0.42 | 0.55 |
| heterochromatin_H3K9me3 | 1000GP | intergenic | -0.90 | -0.97 | -0.83 |
| heterochromatin_H3K9me3 | ADNI | noncoding | 1.59 | 1.50 | 1.69 |
| heterochromatin_H3K9me3 | ADNI | intronic | -1.58 | -1.67 | -1.48 |
| heterochromatin_H3K9me3 | ADNI | intergenic | 2.88 | 2.74 | 3.02 |

**Table S5d3. Depletion simulation results for DHS or enhancer (PlyRS$_{sum\text{-}pleio}$).**

| regclass | dataset | gencompartment | cohensd | cohensd_lower | cohensd_upper |
|---|---|---|---|---|---|
| DHS_hotspot | 1000GP | noncoding | 6.67 | 6.38 | 6.97 |
| DHS_hotspot | 1000GP | intronic | 5.05 | 4.82 | 5.28 |
| DHS_hotspot | 1000GP | intergenic | 4.32 | 4.12 | 4.51 |
| DHS_hotspot | ADNI | noncoding | 4.96 | 4.73 | 5.19 |
| DHS_hotspot | ADNI | intronic | 3.61 | 3.44 | 3.78 |
| DHS_hotspot | ADNI | intergenic | 3.51 | 3.35 | 3.68 |
| DHS_MACS | 1000GP | noncoding | 6.99 | 6.67 | 7.30 |
| DHS_MACS | 1000GP | intronic | 5.44 | 5.19 | 5.69 |
| DHS_MACS | 1000GP | intergenic | 4.19 | 4.00 | 4.39 |
| DHS_MACS | ADNI | noncoding | 5.22 | 4.98 | 5.46 |
| DHS_MACS | ADNI | intronic | 3.56 | 3.39 | 3.73 |
| DHS_MACS | ADNI | intergenic | 3.91 | 3.73 | 4.10 |

| | | | | | |
|---|---|---|---|---|---|
| enhancer_H3K4me1 | 1000GP | noncoding | 4.44 | 4.24 | 4.64 |
| enhancer_H3K4me1 | 1000GP | intronic | 2.99 | 2.84 | 3.13 |
| enhancer_H3K4me1 | 1000GP | intergenic | 3.68 | 3.50 | 3.85 |
| enhancer_H3K4me1 | ADNI | noncoding | 4.04 | 3.85 | 4.22 |
| enhancer_H3K4me1 | ADNI | intronic | 2.89 | 2.75 | 3.03 |
| enhancer_H3K4me1 | ADNI | intergenic | 2.95 | 2.81 | 3.10 |
| transcribed_H3K36me3 | 1000GP | noncoding | -1.90 | -2.00 | -1.80 |
| transcribed_H3K36me3 | 1000GP | intronic | -1.90 | -2.01 | -1.80 |
| transcribed_H3K36me3 | 1000GP | intergenic | 0.21 | 0.15 | 0.27 |
| transcribed_H3K36me3 | ADNI | noncoding | -1.78 | -1.88 | -1.68 |
| transcribed_H3K36me3 | ADNI | intronic | -1.81 | -1.91 | -1.71 |
| transcribed_H3K36me3 | ADNI | intergenic | 0.91 | 0.84 | 0.99 |
| polycomb_H3K27me3 | 1000GP | noncoding | 0.00 | -0.06 | 0.06 |
| polycomb_H3K27me3 | 1000GP | intronic | 1.49 | 1.40 | 1.57 |
| polycomb_H3K27me3 | 1000GP | intergenic | -0.95 | -1.03 | -0.88 |
| polycomb_H3K27me3 | ADNI | noncoding | 1.74 | 1.64 | 1.84 |
| polycomb_H3K27me3 | ADNI | intronic | 1.79 | 1.69 | 1.89 |
| polycomb_H3K27me3 | ADNI | intergenic | 1.00 | 0.92 | 1.07 |
| heterochromatin_H3K9me3 | 1000GP | noncoding | -0.96 | -1.04 | -0.89 |
| heterochromatin_H3K9me3 | 1000GP | intronic | -2.47 | -2.60 | -2.35 |

| | | | | | |
|---|---|---|---|---|---|
| heterochromatin_H3K9me3 | 1000GP | intergenic | 0.26 | 0.20 | 0.33 |
| heterochromatin_H3K9me3 | ADNI | noncoding | -4.72 | -4.94 | -4.51 |
| heterochromatin_H3K9me3 | ADNI | intronic | -2.49 | -2.62 | -2.37 |
| heterochromatin_H3K9me3 | ADNI | intergenic | -3.92 | -4.10 | -3.74 |

**Table S5d4. Depletion simulation results for TAD-loop (binary).**

| regclass | dataset | gencompartment | cohensd | cohensd_lower | cohensd_upper |
|---|---|---|---|---|---|
| TAD-loop | 1000GP | noncoding | 2.63 | 2.50 | 2.76 |
| TAD-loop | 1000GP | intronic | 1.63 | 1.54 | 1.73 |
| TAD-loop | 1000GP | intergenic | 2.10 | 1.99 | 2.21 |
| TAD-loop | ADNI | noncoding | 0.45 | 0.39 | 0.52 |
| TAD-loop | ADNI | intronic | -0.27 | -0.34 | -0.21 |
| TAD-loop | ADNI | intergenic | 0.93 | 0.86 | 1.00 |
| TAD-loop_noCTCFwithin | 1000GP | noncoding | 1.62 | 1.52 | 1.71 |
| TAD-loop_noCTCFwithin | 1000GP | intronic | 0.80 | 0.73 | 0.87 |
| TAD-loop_noCTCFwithin | 1000GP | intergenic | 1.52 | 1.43 | 1.61 |
| TAD-loop_noCTCFwithin | ADNI | noncoding | -0.36 | -0.42 | -0.29 |
| TAD-loop_noCTCFwithin | ADNI | intronic | -0.54 | -0.60 | -0.47 |
| TAD-loop_noCTCFwithin | ADNI | intergenic | 0.06 | 0.00 | 0.12 |
| CTCF-within_TAD-loop | 1000GP | noncoding | 3.76 | 3.59 | 3.94 |

| | | | | | |
|---|---|---|---|---|---|
| CTCF-within_TAD-loop | ADNI | noncoding | 1.37 | 1.28 | 1.45 |

**Table S5d5. Depletion simulation results for TAD-loop (PlyRS$_{max}$).**

| regclass | dataset | gencompartment | cohensd | cohensd_lower | cohensd_upper |
|---|---|---|---|---|---|
| TAD-loop | 1000GP | noncoding | 2.39 | 2.27 | 2.51 |
| TAD-loop | 1000GP | intronic | 1.45 | 1.36 | 1.53 |
| TAD-loop | 1000GP | intergenic | 2.01 | 1.90 | 2.11 |
| TAD-loop | ADNI | noncoding | 0.97 | 0.89 | 1.04 |
| TAD-loop | ADNI | intronic | 0.39 | 0.32 | 0.45 |
| TAD-loop | ADNI | intergenic | 1.00 | 0.93 | 1.08 |
| TAD-loop_noCTCFwithin | 1000GP | noncoding | 1.25 | 1.16 | 1.33 |
| TAD-loop_noCTCFwithin | 1000GP | intronic | 0.48 | 0.41 | 0.54 |
| TAD-loop_noCTCFwithin | 1000GP | intergenic | 1.34 | 1.25 | 1.42 |
| TAD-loop_noCTCFwithin | ADNI | noncoding | -0.06 | -0.12 | 0.01 |
| TAD-loop_noCTCFwithin | ADNI | intronic | -0.34 | -0.40 | -0.27 |
| TAD-loop_noCTCFwithin | ADNI | intergenic | 0.28 | 0.22 | 0.34 |
| CTCF-within_TAD-loop | 1000GP | noncoding | 3.81 | 3.64 | 3.99 |
| CTCF-within_TAD-loop | ADNI | noncoding | 1.20 | 1.12 | 1.28 |

### Note S6

#### Logistic regression and covariates

##### Note S6a

***Logistic regression***

For a regulatory element feature, to test whether PlyRS measure depletion magnitude (SI Note S6b) depends on deletion allele frequency, we use logistic regression on rare (AF<=1%) or common (AF>1%) allele frequency in the presence of genomic covariates. Since the vast majority of deletions in our datasets are rare (~52% of 1000GP deletions and ~76% of ADNI deletions are tripleton or lower in allele frequency), multivariate regression performed directly on allele frequency would create a response variable that does not 'behave well' in terms of its resulting distribution. Therefore, conclusions reached from p-value interpretation would be unreliable. Using logistic regression, we collapse all rare deletions (AF<=1%) into a single class, thereby creating a binary response variable of rare/common, from which multivariate regression can be performed with confidence in the p-value interpretation. We use <=1% as an AF cutoff for rare deletions as it minimizes technical artifacts that might be present from just examining singletons alone (where calling artifacts might predominantly reside in the AFS). Since ADNI deletions are much more rare, on average, than 1000GP deletions and this test collapses all rare deletions into a single class, it may be expected that real associations might appear statistically weaker in the ADNI dataset for this test format.

For genomic covariates of each deletion, we choose average regional measures 50 kilo-bases (kb) upstream of the start coordinate and downstream of the end coordinate (correlation between 50kb sides values and within-deletion values is extremely significant [$p<10^{-16}$]). To calculate SNV nucleotide diversity (pi), we used VCFtools (SI 27) with '--remove-indels' option on the 1000GP data (downloaded from server: http://ftp.1000genomes.ebi.ac.uk/vol1/ftp/release/20130502/) and ADNI data (SI Note S2b and Note S3), examining only sites in the individuals from which the deletion genotypes were derived (2,504 individuals in 1000GP and 752 individuals in ADNI). Recombination rate was taken from the HapMap-derived 'Combined_LD' column for 1000GP and 'CEU_LD' column for ADNI (downloaded from: http://www.well.ox.ac.uk/~anjali/AAmap/). We chose the HapMap data since it covered more of the genome than the other measures. For distance to the nearest transcription start site, we used coordinate information downloaded from Ensembl Biomart (http://grch37.ensembl.org/biomart/martview/ with dataset: ‘Human genes (GRCh37.p13)’). The Ensembl data was used instead of UCSC data because the Ensembl data appeared to contain more transcripts. GC content proportion is calculated directly from the GRCh37/hg19 version of the human reference genome. The reference genome file used was human_g1k_v37.fasta (downloaded from the 1000 Genomes FTP server (ftp://ftp.1000genomes.ebi.ac.uk/vol1/ftp//technical/reference/human_g1k_v37.fasta.gz).

#### Note S6b

***Depletion Magnitude Calculation***

To get meaningful odds ratio interpretation of magnitude of depletion in logistic regression tests, we use the ratio of proportional difference between real deletions and length-matched simulations (sim) (calculated as: [$PlyRS_{measure}$ - average sim $PlyRS_{measure}$] / average sim $PlyRS_{measure}$) for chromatin accessibility and histone modification annotations. This means that a real deletion for which no regulatory overlap occurs will be considered to be 100% depleted (-1.0) of PlyRS measure in relation to the deletion simulation average of the PlyRS measure. We use the raw difference between real deletions and simulations ($PlyRS_{measure}$ - average sim $PlyRS_{measure}$) for TAD-loop and CTCF annotations. We don't use raw difference for chromatin accessibility and histone modification features because otherwise we would length-biasing the values since deletions can potentially overlap more than one of these regulatory loci, however because of the length dynamics (SI Note S5c), deletions can only overlap at most one TAD-loop boundary. The odds ratio in these tests means that for a one unit change in the difference (either proportional or raw) compared to simulations, with all other covariates held steady, there is an odds increase or decrease (with 1 as baseline) of the deletion set depletion magnitude being positively associated with allele frequency (i.e. deletions more depleted from simulation average are more likely to be common). Confidence intervals on the odds ratio are calculated as profile likelihood based confidence intervals, as we are not able to depend on the assumption of normality for the estimator.

#### Note S6c

***Logistic regression results***

**Table S6c1. Logistic regression results for DHS, enhancer, transcribed, polycomb-repressed, or heterochromatin ($PlyRS_{sum}$).**

| regclass | dataset | gencompartment | OR_estimate | OR_estimate_lower | OR_estimate_upper |
|---|---|---|---|---|---|
| DHS_hotspot | 1000GP | noncoding | 1.07 | 1.04 | 1.11 |
| DHS_hotspot | 1000GP | intronic | 1.09 | 1.04 | 1.14 |
| DHS_hotspot | 1000GP | intergenic | 1.06 | 1.02 | 1.10 |
| DHS_hotspot | ADNI | noncoding | 1.11 | 1.01 | 1.23 |
| DHS_hotspot | ADNI | intronic | 1.09 | 0.94 | 1.29 |
| DHS_hotspot | ADNI | intergenic | 1.12 | 1.00 | 1.28 |
| DHS_MACS | 1000GP | noncoding | 1.09 | 1.05 | 1.13 |
| DHS_MACS | 1000GP | intronic | 1.11 | 1.06 | 1.18 |
| DHS_MACS | 1000GP | intergenic | 1.07 | 1.03 | 1.12 |
| DHS_MACS | ADNI | noncoding | 1.17 | 1.05 | 1.31 |
| DHS_MACS | ADNI | intronic | 1.08 | 0.92 | 1.29 |

| | | | | | |
|---|---|---|---|---|---|
| DHS_MACS | ADNI | intergenic | 1.23 | 1.07 | 1.43 |
| enhancer_H3K4me1 | 1000GP | noncoding | 1.06 | 1.02 | 1.10 |
| enhancer_H3K4me1 | 1000GP | intronic | 1.11 | 1.05 | 1.18 |
| enhancer_H3K4me1 | 1000GP | intergenic | 1.04 | 1.00 | 1.08 |
| enhancer_H3K4me1 | ADNI | noncoding | 1.09 | 0.99 | 1.20 |
| enhancer_H3K4me1 | ADNI | intronic | 0.96 | 0.81 | 1.14 |
| enhancer_H3K4me1 | ADNI | intergenic | 1.16 | 1.04 | 1.31 |
| transcribed_H3K36me3 | 1000GP | noncoding | 1.02 | 1.00 | 1.04 |
| transcribed_H3K36me3 | 1000GP | intronic | 1.07 | 1.02 | 1.12 |
| transcribed_H3K36me3 | 1000GP | intergenic | 1.01 | 1.00 | 1.03 |
| transcribed_H3K36me3 | ADNI | noncoding | 1.00 | 0.97 | 1.03 |
| transcribed_H3K36me3 | ADNI | intronic | 1.07 | 0.96 | 1.22 |
| transcribed_H3K36me3 | ADNI | intergenic | 0.99 | 0.97 | 1.02 |
| polycomb_H3K27me3 | 1000GP | noncoding | 1.00 | 0.98 | 1.03 |
| polycomb_H3K27me3 | 1000GP | intronic | 0.99 | 0.95 | 1.03 |
| polycomb_H3K27me3 | 1000GP | intergenic | 1.03 | 0.99 | 1.07 |
| polycomb_H3K27me3 | ADNI | noncoding | 1.02 | 0.94 | 1.12 |
| polycomb_H3K27me3 | ADNI | intronic | 0.94 | 0.84 | 1.06 |
| polycomb_H3K27me3 | ADNI | intergenic | 1.13 | 1.00 | 1.28 |
| heterochromatin_H3K9me3 | 1000GP | noncoding | 1.03 | 1.00 | 1.06 |
| heterochromatin_H3K9me3 | 1000GP | intronic | 1.04 | 1.00 | 1.08 |

| | | | | | |
|---|---|---|---|---|---|
| heterochromatin_H3K9me3 | 1000GP | intergenic | 1.02 | 0.98 | 1.07 |
| heterochromatin_H3K9me3 | ADNI | noncoding | 1.06 | 0.98 | 1.15 |
| heterochromatin_H3K9me3 | ADNI | intronic | 1.00 | 0.90 | 1.11 |
| heterochromatin_H3K9me3 | ADNI | intergenic | 1.14 | 1.02 | 1.29 |

**Table S6c2. Logistic regression results for DHS or enhancer ($PlyRS_{sum\text{-}mono}$).**

| regclass | dataset | gencompartment | OR_estimate | OR_estimate_lower | OR_estimate_upper |
|---|---|---|---|---|---|
| DHS_hotspot | 1000GP | noncoding | 1.02 | 1.00 | 1.04 |
| DHS_hotspot | 1000GP | intronic | 1.02 | 0.99 | 1.05 |
| DHS_hotspot | 1000GP | intergenic | 1.02 | 0.99 | 1.05 |
| DHS_hotspot | ADNI | noncoding | 1.04 | 0.97 | 1.13 |
| DHS_hotspot | ADNI | intronic | 1.03 | 0.92 | 1.17 |
| DHS_hotspot | ADNI | intergenic | 1.05 | 0.95 | 1.17 |
| DHS_MACS | 1000GP | noncoding | 1.04 | 1.01 | 1.07 |
| DHS_MACS | 1000GP | intronic | 1.04 | 1.00 | 1.09 |
| DHS_MACS | 1000GP | intergenic | 1.04 | 1.00 | 1.07 |
| DHS_MACS | ADNI | noncoding | 1.13 | 1.02 | 1.27 |
| DHS_MACS | ADNI | intronic | 1.02 | 0.87 | 1.20 |
| DHS_MACS | ADNI | intergenic | 1.23 | 1.06 | 1.45 |
| enhancer_H3K4me1 | 1000GP | noncoding | 1.02 | 0.99 | 1.04 |
| enhancer_H3K4me1 | 1000GP | intronic | 1.00 | 0.97 | 1.04 |
| enhancer_H3K4me1 | 1000GP | intergenic | 1.03 | 1.00 | 1.06 |
| enhancer_H3K4me1 | ADNI | noncoding | 1.08 | 1.00 | 1.18 |
| enhancer_H3K4me1 | ADNI | intronic | 1.01 | 0.89 | 1.15 |
| enhancer_H3K4me1 | ADNI | intergenic | 1.14 | 1.02 | 1.28 |

| | | | | | |
|---|---|---|---|---|---|
| transcribed_H3K36me3 | 1000GP | noncoding | 1.01 | 1.00 | 1.02 |
| transcribed_H3K36me3 | 1000GP | intronic | 1.01 | 0.97 | 1.04 |
| transcribed_H3K36me3 | 1000GP | intergenic | 1.01 | 0.99 | 1.02 |
| transcribed_H3K36me3 | ADNI | noncoding | 1.00 | 0.96 | 1.04 |
| transcribed_H3K36me3 | ADNI | intronic | 1.01 | 0.91 | 1.14 |
| transcribed_H3K36me3 | ADNI | intergenic | 0.99 | 0.96 | 1.04 |
| polycomb_H3K27me3 | 1000GP | noncoding | 1.00 | 0.98 | 1.02 |
| polycomb_H3K27me3 | 1000GP | intronic | 0.99 | 0.96 | 1.02 |
| polycomb_H3K27me3 | 1000GP | intergenic | 1.02 | 0.99 | 1.05 |
| polycomb_H3K27me3 | ADNI | noncoding | 1.00 | 0.93 | 1.08 |
| polycomb_H3K27me3 | ADNI | intronic | 0.97 | 0.88 | 1.09 |
| polycomb_H3K27me3 | ADNI | intergenic | 1.03 | 0.93 | 1.15 |
| heterochromatin_H3K9me3 | 1000GP | noncoding | 1.02 | 1.00 | 1.04 |
| heterochromatin_H3K9me3 | 1000GP | intronic | 1.03 | 1.00 | 1.07 |
| heterochromatin_H3K9me3 | 1000GP | intergenic | 1.00 | 0.97 | 1.04 |
| heterochromatin_H3K9me3 | ADNI | noncoding | 1.09 | 0.99 | 1.20 |
| heterochromatin_H3K9me3 | ADNI | intronic | 1.11 | 0.98 | 1.28 |
| heterochromatin_H3K9me3 | ADNI | intergenic | 1.06 | 0.93 | 1.23 |

**Table S6c3. Logistic regression results for DHS or enhancer (PlyRS$_{sum\text{-}pleio}$).**

| regclass | dataset | gencompartment | OR_estimate | OR_estimate_lower | OR_estimate_upper |
|---|---|---|---|---|---|

| | | | | | |
|---|---|---|---|---|---|
| DHS_hotspot | 1000GP | noncoding | 1.04 | 1.02 | 1.06 |
| DHS_hotspot | 1000GP | intronic | 1.06 | 1.03 | 1.10 |
| DHS_hotspot | 1000GP | intergenic | 1.02 | 1.00 | 1.05 |
| DHS_hotspot | ADNI | noncoding | 1.07 | 1.00 | 1.16 |
| DHS_hotspot | ADNI | intronic | 1.05 | 0.96 | 1.19 |
| DHS_hotspot | ADNI | intergenic | 1.08 | 0.99 | 1.20 |
| DHS_MACS | 1000GP | noncoding | 1.04 | 1.02 | 1.06 |
| DHS_MACS | 1000GP | intronic | 1.07 | 1.04 | 1.12 |
| DHS_MACS | 1000GP | intergenic | 1.02 | 1.00 | 1.05 |
| DHS_MACS | ADNI | noncoding | 1.07 | 1.01 | 1.16 |
| DHS_MACS | ADNI | intronic | 1.06 | 0.97 | 1.21 |
| DHS_MACS | ADNI | intergenic | 1.08 | 1.00 | 1.21 |
| enhancer_H3K4me1 | 1000GP | noncoding | 1.03 | 1.01 | 1.05 |
| enhancer_H3K4me1 | 1000GP | intronic | 1.06 | 1.02 | 1.10 |
| enhancer_H3K4me1 | 1000GP | intergenic | 1.02 | 0.99 | 1.04 |
| enhancer_H3K4me1 | ADNI | noncoding | 1.04 | 0.99 | 1.11 |
| enhancer_H3K4me1 | ADNI | intronic | 0.99 | 0.90 | 1.11 |
| enhancer_H3K4me1 | ADNI | intergenic | 1.07 | 1.00 | 1.16 |
| transcribed_H3K36me3 | 1000GP | noncoding | 1.01 | 1.00 | 1.02 |
| transcribed_H3K36me3 | 1000GP | intronic | 1.05 | 1.02 | 1.08 |
| transcribed_H3K36me3 | 1000GP | intergenic | 1.00 | 1.00 | 1.01 |
| transcribed_H3K36me3 | ADNI | noncoding | 1.00 | 0.99 | 1.01 |
| transcribed_H3K36me3 | ADNI | intronic | 1.08 | 1.00 | 1.18 |
| transcribed_H3K36me3 | ADNI | intergenic | 1.00 | 0.99 | 1.01 |

| | | | | | |
|---|---|---|---|---|---|
| polycomb_H3K27me3 | 1000GP | noncoding | 1.00 | 0.99 | 1.02 |
| polycomb_H3K27me3 | 1000GP | intronic | 1.00 | 0.99 | 1.02 |
| polycomb_H3K27me3 | 1000GP | intergenic | 1.01 | 0.99 | 1.04 |
| polycomb_H3K27me3 | ADNI | noncoding | 1.02 | 0.98 | 1.07 |
| polycomb_H3K27me3 | ADNI | intronic | 0.99 | 0.96 | 1.05 |
| polycomb_H3K27me3 | ADNI | intergenic | 1.07 | 1.00 | 1.17 |
| heterochromatin_H3K9me3 | 1000GP | noncoding | 1.01 | 1.00 | 1.03 |
| heterochromatin_H3K9me3 | 1000GP | intronic | 1.01 | 1.00 | 1.03 |
| heterochromatin_H3K9me3 | 1000GP | intergenic | 1.02 | 0.99 | 1.04 |
| heterochromatin_H3K9me3 | ADNI | noncoding | 1.03 | 1.00 | 1.07 |
| heterochromatin_H3K9me3 | ADNI | intronic | 1.00 | 0.96 | 1.05 |
| heterochromatin_H3K9me3 | ADNI | intergenic | 1.07 | 1.02 | 1.15 |

**Table S6c4. Logistic regression results for TAD-loop (binary).**

| regclass | dataset | gencompartment | OR_estimate | OR_estimate_lower | OR_estimate_upper |
|---|---|---|---|---|---|
| TAD-loop | 1000GP | noncoding | 1.19 | 1.03 | 1.39 |
| TAD-loop | 1000GP | intronic | 1.13 | 0.92 | 1.40 |
| TAD-loop | 1000GP | intergenic | 1.28 | 1.03 | 1.62 |
| TAD-loop | ADNI | noncoding | 1.66 | 1.15 | 2.49 |
| TAD-loop | ADNI | intronic | 1.64 | 0.96 | 2.98 |
| TAD-loop | ADNI | intergenic | 1.71 | 1.02 | 3.04 |
| TAD-loop_noCTCFwithin | 1000GP | noncoding | 1.15 | 0.98 | 1.34 |

| | | | | | |
|---|---|---|---|---|---|
| TAD-loop_noCTCFwithin | 1000GP | intronic | 1.12 | 0.91 | 1.39 |
| TAD-loop_noCTCFwithin | 1000GP | intergenic | 1.19 | 0.95 | 1.51 |
| TAD-loop_noCTCFwithin | ADNI | noncoding | 1.42 | 0.97 | 2.15 |
| TAD-loop_noCTCFwithin | ADNI | intronic | 1.37 | 0.80 | 2.49 |
| TAD-loop_noCTCFwithin | ADNI | intergenic | 1.50 | 0.89 | 2.73 |
| CTCF-within_TAD-loop | 1000GP | noncoding | 2.70 | 1.35 | 6.37 |
| CTCF-within_TAD-loop | ADNI | noncoding | 7.67 | 1.76 | 108.20 |

**Table S6c5. Logistic regression results for TAD-loop (PlyRS$_{max}$).**

| regclass | dataset | gencompartment | OR_estimate | OR_estimate_lower | OR_estimate_upper |
|---|---|---|---|---|---|
| TAD-loop | 1000GP | noncoding | 1.46 | 1.01 | 2.17 |
| TAD-loop | 1000GP | intronic | 1.20 | 0.72 | 2.07 |
| TAD-loop | 1000GP | intergenic | 1.84 | 1.07 | 3.31 |
| TAD-loop | ADNI | noncoding | 4.13 | 1.45 | 14.24 |
| TAD-loop | ADNI | intronic | 3.93 | 0.82 | 28.51 |
| TAD-loop | ADNI | intergenic | 4.44 | 1.16 | 24.13 |
| TAD-loop_noCTCFwithin | 1000GP | noncoding | 1.28 | 0.87 | 1.93 |
| TAD-loop_noCTCFwithin | 1000GP | intronic | 1.18 | 0.70 | 2.08 |
| TAD-loop_noCTCFwithin | 1000GP | intergenic | 1.44 | 0.82 | 2.62 |

| | | | | | |
|---|---|---|---|---|---|
| TAD-loop_noCTCFwithin | ADNI | noncoding | 3.02 | 1.02 | 10.93 |
| TAD-loop_noCTCFwithin | ADNI | intronic | 2.19 | 0.45 | 15.79 |
| TAD-loop_noCTCFwithin | ADNI | intergenic | 4.04 | 0.95 | 25.41 |
| CTCF-within_TAD-loop | 1000GP | noncoding | 36.80 | 3.49 | 1,277.08 |
| CTCF-within_TAD-loop | ADNI | noncoding | 30.11 | 1.27 | 8,491.69 |

Figure S1

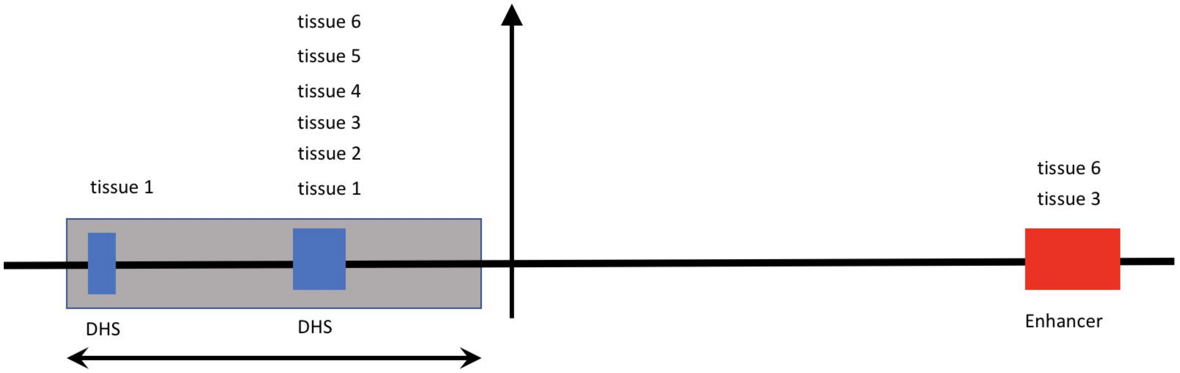


**Fig. S1. Purifying selection against deletions can operate along two 'axes' at a genomic locus.** A deletion can overlap one or more regulatory elements (here shaded grey, overlapping two DHS sites, but missing an enhancer site), thus removing putative genomic function along a horizontal axis. The regulatory element(s) overlapped can have tissue/cell-type-specific function (as the DHS on the left side) or cellularly pleiotropic (i.e. more than one tissue/cell-type) function (as the DHS on the right side), thus removing putative genomic function along a vertical axis.

Figure S2

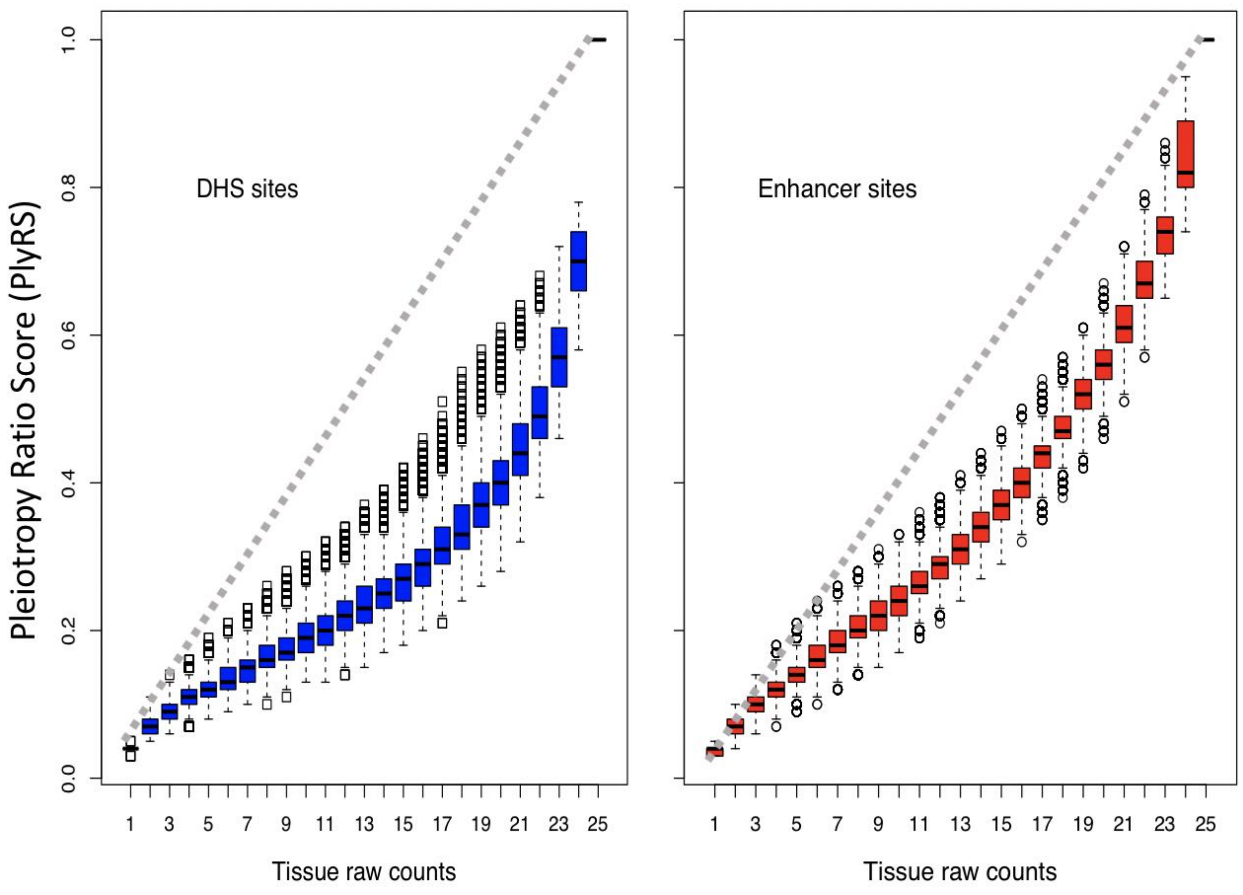


**Fig. S2. Comparison of PlyRS to raw tissue counts.** PlyRS can range from 0, representing no regulatory activity in any tissue/cell-type, to 1, representing regulatory activity in all tissues/cell-types analyzed. Because PlyRS accounts for the positive regulatory activity correlation between tissues and cell-types analyzed, PlyRS (y-axis) will always fall at, or below, the diagonal versus a simple raw count (x-axis). Each regulatory feature will display a different PlyRS distribution (e.g. the enhancer_H3K4me1 feature [right] has a PlyRS distribution that lies closer to the diagonal than for the DHS_hotspot feature [left]), based on the activity covariance of the tissues/cell-types that we analyzed (SI Note S4) of that regulatory feature.

Figure S3

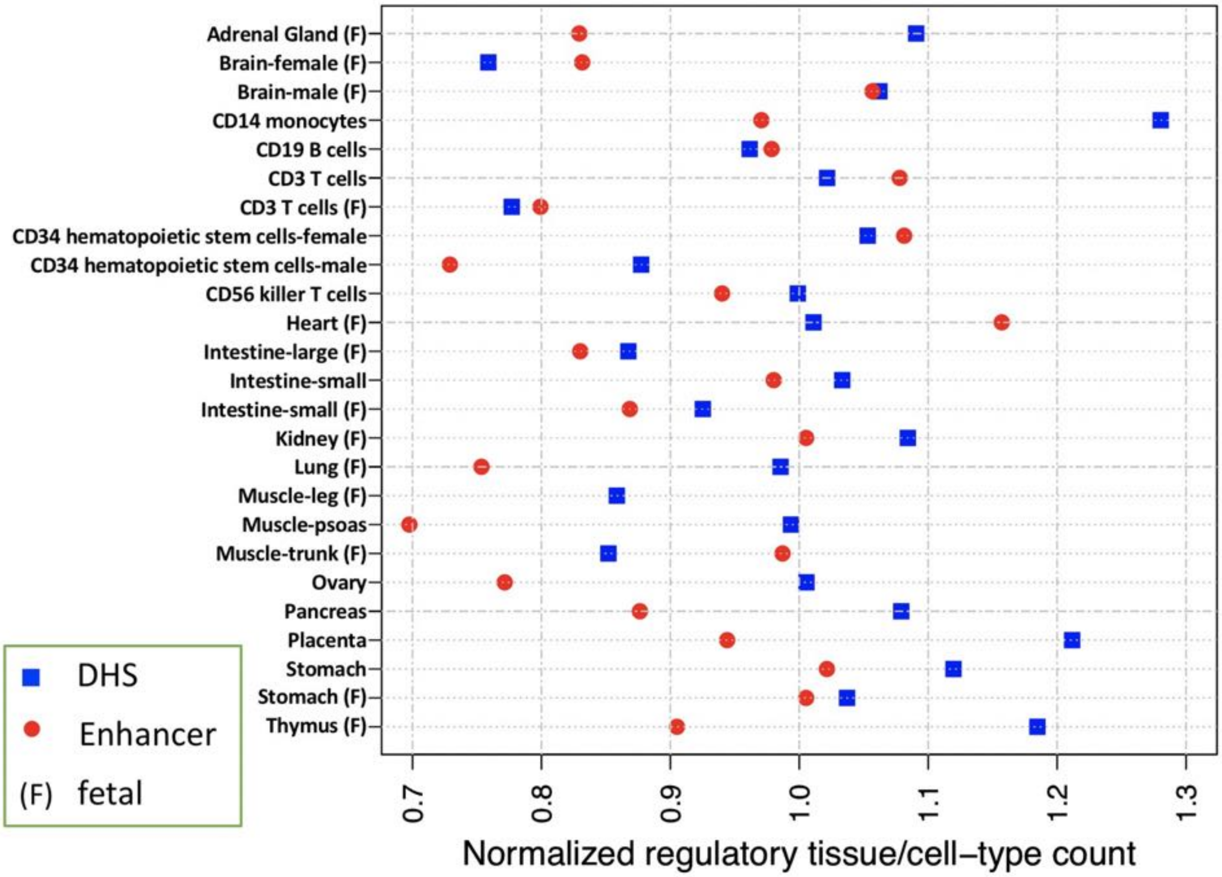


**Fig. S3. Comparison of PlyRS for tissue/cell-type-specific counts.** PlyRS accounts for the regulatory activity correlation between tissues and cell-types analyzed, and will consequently up-weight tissues/cell-types that genome-wide have relatively rare activity and down-weight tissues/cell-types with relatively common activity. When cellular activity is found to be tissue/cell-type-specific, each tissue/cell-type will have a PlyRS corresponding to it that may be below, at, or above a normalized simple count of 1. Because of the highly-correlated nature of DHS_hotspot (blue) sites and enhancer_H3K4me1 (red) sites among the tissues/cell-types that we analyzed, most tissues and cell-types fall below a normalized count of 1 when appearing at a locus as tissue/cell-type-specific.

Figure S4

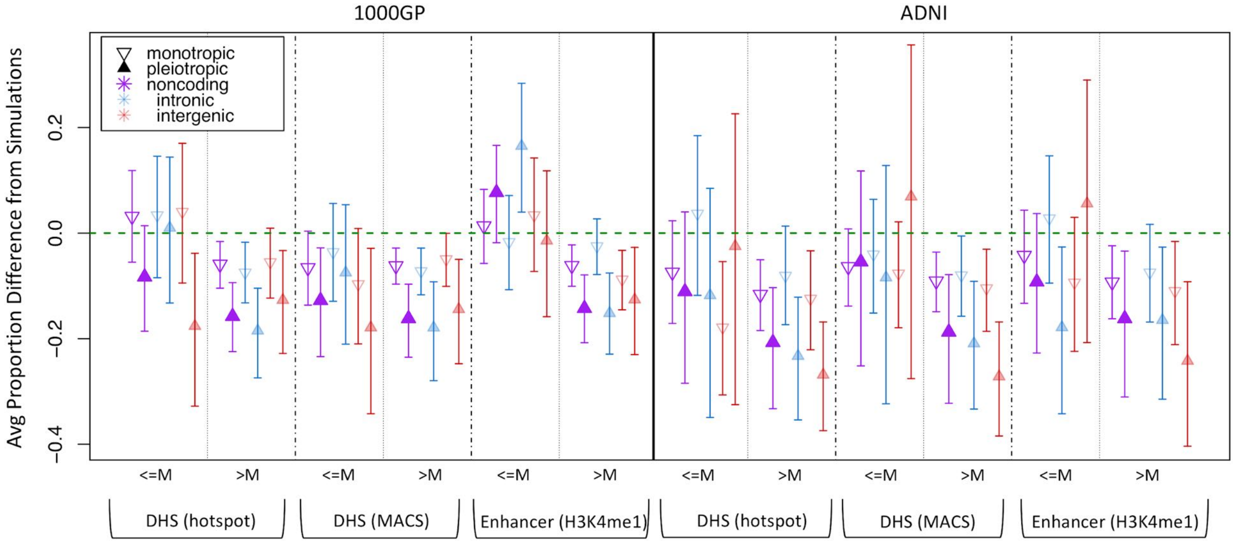


**Fig. S4. Selective burden in deletions based on length.** We calculated the depletion magnitude of each deletion (*SI Appendix*, *Note S6b*) in both the 1000GP dataset (left half) and the ADNI dataset (right half) in terms of $PlyRS_{sum\text{-}mono}$ (monotropic) and $PlyRS_{sum\text{-}pleio}$ (pleiotropic). We separated deletions from each dataset into those that had a length of less than or equal to the median (M) length or greater than the median length of the deletion subset. For regulatory annotations (listed on the bottom) we plot the average $PlyRS_{sum\text{-}mono}$ (or $PlyRS_{sum\text{-}pleio}$) depletion magnitude (proportional difference) for each deletion subset and generate 95% confidence intervals from bootstrapping (100 trials). A more negative value for the average proportional difference indicates a larger average depletion magnitude of real deletions compared to simulation. There appears to be a qualitative trend (not statistically significant) of more negative average proportional difference of longer deletions (>M) compared to shorter deletions (<=M), especially for pleiotropic sets.

Table S1

**1000GP Filtered Deletion Callset characteristics**

| | |
|---|---|
| Number of Deletions | 12,013 (100%) |
| Number of Intronic Deletions | 5,896 (49.1%) |
| Number of Intergenic Deletions | 6,117 (50.9%) |
| Singleton AF | 4,362 (36.3%) |
| Doubleton AF | 1,241 (10.3%) |
| Tripleton AF | 617 (5.1%) |
| >1% AF | 2,165 (18.0%) |
| Average Length | 1,437 bp |
| Median Length | 629 bp |
| Minimum Length | 50 bp |
| Maximum Length | 22,648 bp |
| Average Length Singleton AF | 1,400 bp |
| Median Length Singleton AF | 448 bp |
| Average Length >1% AF | 979 bp |
| Median Length >1% AF | 355 bp |

#### Table S2

**ADNI Filtered Deletion Callset characteristics**

| | |
|---|---|
| Number of Deletions | 3,306 (100%) |
| Number of Intronic Deletions | 1,459 (44.1%) |
| Number of Intergenic Deletions | 1,847 (55.9%) |
| Singleton AF | 2,007 (60.7%) |
| Doubleton AF | 327 (9.9%) |
| Tripleton AF | 163 (4.9%) |
| >1% AF | 395 (11.9%) |
| Average Length | 2,722 bp |
| Median Length | 1,893 bp |
| Minimum Length | 440 bp |
| Maximum Length | 23,344 bp |
| Average Length Singleton AF | 2,760 bp |
| Median Length Singleton AF | 1,924 bp |
| Average Length >1% AF | 3,072 bp |
| Median Length >1% AF | 2,297 bp |

#### Table S3

**Tissues and cell-types analyzed from REC**

| Tissue/Cell-type Description | Consolidated Epigenome ID | Fetal (F) or Adult (A) |
|---|---|---|
| Primary monocytes from peripheral blood | E029 | A |
| Primary B cells from peripheral blood | E032 | A |
| Primary T cells from cord blood | E033 | F |
| Primary T cells from peripheral blood | E034 | A |
| Primary Natural Killer cells from peripheral blood | E046 | A |
| Primary hematopoietic stem cells G-CSF-mobilized Female | E050 | F |
| Primary hematopoietic stem cells G-CSF-mobilized Male | E051 | A |
| Fetal Adrenal Gland | E080 | F |
| Fetal Brain Male | E081 | F |
| Fetal Brain Female | E082 | F |
| Fetal Heart | E083 | F |
| Fetal Intestine Large | E084 | F |
| Fetal Intestine Small | E085 | F |
| Fetal Kidney | E086 | F |
| Fetal Lung | E088 | F |
| Fetal Muscle Trunk | E089 | F |
| Fetal Muscle Leg | E090 | F |
| Placenta | E091 | F |
| Fetal Stomach | E092 | F |
| Fetal Thymus | E093 | F |
| Gastric | E094 | A |
| Ovary | E097 | A |
| Pancreas | E098 | A |
| Psoas Muscle | E100 | A |
| Small Intestine | E109 | A |
